# Stereoselective Degradation of Diacylglycerol Kinases Potentiate T cell Activation and Tumor Cell Cytotoxicity

**DOI:** 10.64898/2025.12.09.692983

**Authors:** Minhaj Shaikh, Surya Pravo Mookherjee, Claire C. Weckerly, Adam H. Libby, Aizhen Xiao, Yunge Zhao, Sagar D. Vaidya, AeRyon Kim, Zhihong Li, Madeleine L. Ware, Michelle Marants, Olivia L. Murtagh, Wesley J. Wolfe, Timothy NJ Bullock, Benjamin W. Purow, Gerald R. V. Hammond, Ku-Lung Hsu

## Abstract

Stereoselective recognition is a powerful means to differentiate selective versus non-specific activity of small molecules in complex biological systems. Here, we disclose stereochemically defined, sulfonyl-triazole inhibitors of the lipid enzyme diacylglycerol kinase-alpha (DGKα), a key metabolic checkpoint for T cell effector function. Acute treatment with the covalent DGKα inhibitor AHL-7160 recruited endogenous DGKα to the plasma membrane in a stereoselective and isozyme-specific manner. The membrane translocation activity of AHL-7160 correlated with blockade of cellular phosphatidic acid production and potentiation of primary T cell-mediated killing of a glioblastoma cell line. Quantitative chemoproteomics revealed Y669 and K411 as sites of AHL-7160 modification on endogenous DGKα in cells. Extended treatments resulted in proteasome-dependent and proteome-wide selective degradation of DGKα in T cells. Collectively, these findings establish covalent DGKα ligands as potent molecular glues with translational potential in immunotherapy.

## INTRODUCTION

Immune checkpoint blockade has transformed cancer treatment by harnessing the body’s immune system to recognize and kill tumor cells^1–3^. However, tumor infiltrating T cells (TILs) often present a dysfunctional phenotype characterized by poor proliferation, susceptibility to apoptosis, and compromised cytotoxic functions, which is thought to be mediated by expression of immune checkpoint molecules that attenuate signaling^4^. Checkpoint inhibitors targeting inhibitory receptors (CTLA-4^5,6^, PD-1^7,8^, PD-L1^9,10^) release the ‘brakes’ on T cells to unleash effective and durable antitumor responses in melanoma, non-small cell lung cancer and renal cell carcinoma^2,10^. Patient response rates, however, are variable and many develop resistance, highlighting the need for new immunotherapy strategies.

T cell receptor (TCR) signaling is mediated by secondary messengers including diacylglycerols (DAGs) that act as signals to alter subcellular localization and activation of key proteins (e.g. MAPK^11^ and PKC^12^) that are essential for T cell activation^13^. Diacylglycerol kinase-alpha (DGKα) and –zeta (DGKζ) are important negative regulators of TCR signaling by phosphorylating the secondary messenger DAG to terminate its signaling activity^14^ (Figure S1A). Excessive DGK activity and thus attenuated DAG signaling has been linked to defective T cell function^14–27^. The product of DGK metabolism, phosphatidic acid (PA) is also involved in T cell activation^28^. By regulating cellular DAG and PA levels, DGKα and DGKζ function as key metabolic checkpoints for controlling the signaling threshold of T cell priming^29^ (Figure S1A). Selective inhibitors for one or both enzymes could overcome immunosuppression in tumors by restoring T cell effector functions.

DGKα and DGKζ are members of a larger family of lipid kinases that consists of 10 known human isozymes^18,30,31^. Developing DGK isozyme-selective inhibitors remains challenging due in part to a limited number of chemotypes^32,33^ but has recently gained momentum with the release of newer scaffolds including quinolone, naphthyridinone and pyridopyrimidinones for targeting DGKα and/or DGKζ^29,34–37^ (Figure S1C). These inhibitors showed potent biochemical, cell biological and *in vivo* activity as monotherapy or in combination with immune checkpoint blockade and several compounds have entered clinical trials (NCT05407675, NCT05858164, NCT05614102). While these next-generation inhibitors show considerable promise, they also raise important questions about their selectivity across the DGK superfamily and the broader proteome in cells, presenting opportunities for further investigation. The limited structural information available for full-length DGKs along with inconclusive evidence of target engagement with native enzymes — including direct small molecule interaction sites — highlights the potential to develop new modalities that can guide the future design of DGK inhibitors.

Here, we developed stereochemically defined sulfonyl-triazole inhibitors that covalently engage native DGKα in a stereoselective manner. Cellular treatment with the DGKα chiral ligands induced membrane translocation of endogenous DGKα but no other isozymes from each DGK subtype tested. T cells treated with DGKα chiral ligands exhibited rapid blockade of PA production, potentiation of TCR signaling, and cytotoxic activity towards a poorly immunogenic glioblastoma cell line. Finally, DGKα chiral ligands act as molecular glues to selectively induce proteasome-dependent and stereoselective degradation of DGKα in T cells.

## RESULTS

### Developing stereochemically defined covalent DGKα inhibitors

Sulfonyl-triazoles (SuTEx) are electrophilic molecules used for covalent targeting of functional tyrosine and lysine residues on diverse proteins including kinases^38^. Our rationale for developing a covalent DGK inhibitor include potential access to allosteric sites and facile target engagement and selectivity assessment in cells. Recent reports disclosed cyano-substituted naphthyridinone compounds as potent DGKα and DGKζ inhibitors capable of potentiating antitumor activity of T cells^29,34–36^. These compounds consist of three principal regions with a cyanonaphthyridinone headgroup (A-ring), dialkylated piperazine linker (B-ring) and a diversity region (C-ring). The SuTEx warhead was installed in the functional group-tolerant C-ring and configured so that the A/B-ring remains bound to protein after covalent reaction (AHL-7160, Figure 1A and B).

**Figure 1.**
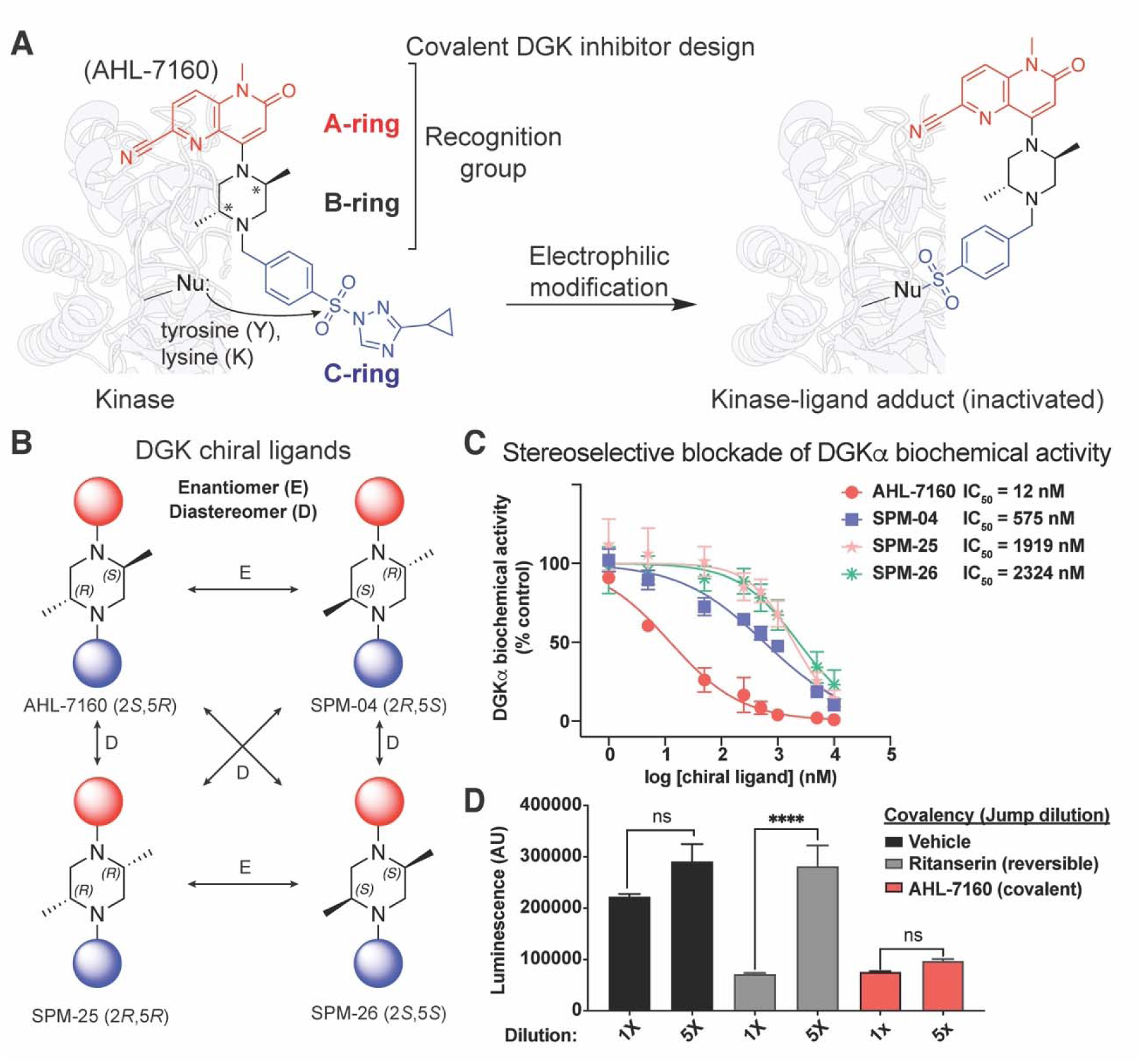
Development of a stereoselective covalent DGKα inhibitor. (A) Schematic showing the design of the covalent DGKα inhibitor AHL-7160. Nucleophilic attack of the sulfonyl-triazole electrophile on AHL-7160 by a lysine and/or tyrosine residue results in covalent protein modification. (B) Generation of a stereochemically-defined library for evaluating stereoselective activity of DGKα covalent inhibitors. (C) Micelle-based biochemical assay for assessing inhibitory activity of DGKα chiral ligands against recombinant hDGKα-HEK293T soluble proteomes. (D) Jump-dilution experiments support covalent binding mechanism of AHL-7160 as evidenced by sustained inhibition upon 5-fold (5X) dilution of reaction mixture. The reversible inhibitor control ritanserin showed the expected loss of inhibitory activity upon 5X dilution. Data shown are representative of n = 2-3 biological replicates.

We used a reported micelle-based DAG phosphorylation biochemical assay^39^ to test AHL-7160 activity. Pretreatment of recombinant human DGKα-HEK293T proteomes with AHL-7160 resulted in concentration-dependent inhibition of DGKα with nanomolar potency (IC_50_ of ∼12 nM, Figure 1C). Next, AHL-7160 was evaluated for DGKζ inhibitory activity since cyanonaphthyridinones were reported as dual-DGKα/DGKζ inhibitors^34,35^. AHL-7160 showed substantially reduced activity (>100-fold) against recombinant DGKζ in the substrate assay (Figure S2A). A ‘reverse-SuTEx’ analog with the A/B-ring positioned on the triazole leaving group substantially reduced activity (SMS-55, Figure S2B).

The stereocenters in the B-ring of AHL-7160 (2*S*,5*R*) provided an opportunity for testing stereoselectivity of DGKα inhibitory activity. We synthesized and tested the enantiomer SPM-04 (2*R*, 5*S*) and observed a greater than 50-fold reduction in activity compared with AHL-7160 (IC_50_ = 575 vs 12 nM for SPM-04 compared with AHL-7160, respectively). The diastereomers SPM-25 (2*R*,5*R*) and –26 (2*S*,5*S*) showed substantially reduced (>160-fold) potency compared with AHL-7160 and SPM-04 (Figure 1B and C). Covalency of AHL-7160 was supported by jump dilution assays where inactivation was retained in contrast with the near complete loss of inhibitory activity using a reversible DGKα inhibitor (Figure 1D). We also tested sulfonic acid and sulfonamide analogs of AHL-7160 and found these reversible counterparts were inactive against DGKα (Figure S2C).

In summary, we show the SuTEx electrophile can be retrofitted into reversible inhibitor scaffolds to develop stereoselective DGKα ligands for guiding downstream biological evaluation.

### AHL-7160 stereoselectively and isozyme specifically recruits DGKα to the plasma membrane

Reversible dual-DGKα/ζ inhibitors, including the advanced compound BMS-986408^36^, were reported to translocate DGKα and DGKζ to the plasma membrane independent of phosphorylation or calcium signals^35,36^. The translocation effects were proposed based on increased binding of DGKα to lipid photoaffinity probes or immunofluorescence (IF) detection of translocated recombinant YFP-DGK proteins^35,36^. Here, we evaluated the ability of AHL-7160 to directly recruit endogenous DGKs to membranes to assess DGK family-wide selectivity of covalent chiral ligands.

We used CRISPR-Cas9 to knock-in the 11^th^ strand of the NeonGreen beta-barrel (NG2^40^) into the N-terminus of DGKα. Upon DGKα expression, the NG211 tag reconstitutes the full NG fluorophore in HEK293A cells stably expressing NG2 strands 1-10. We confirmed successful editing by PCR and sequencing (Figure 2A). Treatment with AHL-7160 resulted in rapid (2-12 min timescale) and concentration dependent recruitment of endogenous NG-DGKα to the plasma membrane as quantified by TIRF microscopy-mediated detection of DGKα particles (AHL-7160, EC_50_ = 39 nM). Treatment with the enantiomer SPM-04 resulted in a ∼50-fold loss in potency of translocation, which matched the stereoselective activity observed in biochemical assays (SPM-04, EC_50_ = 1881 nM; Figure 2B-D, S3).

**Figure 2.**
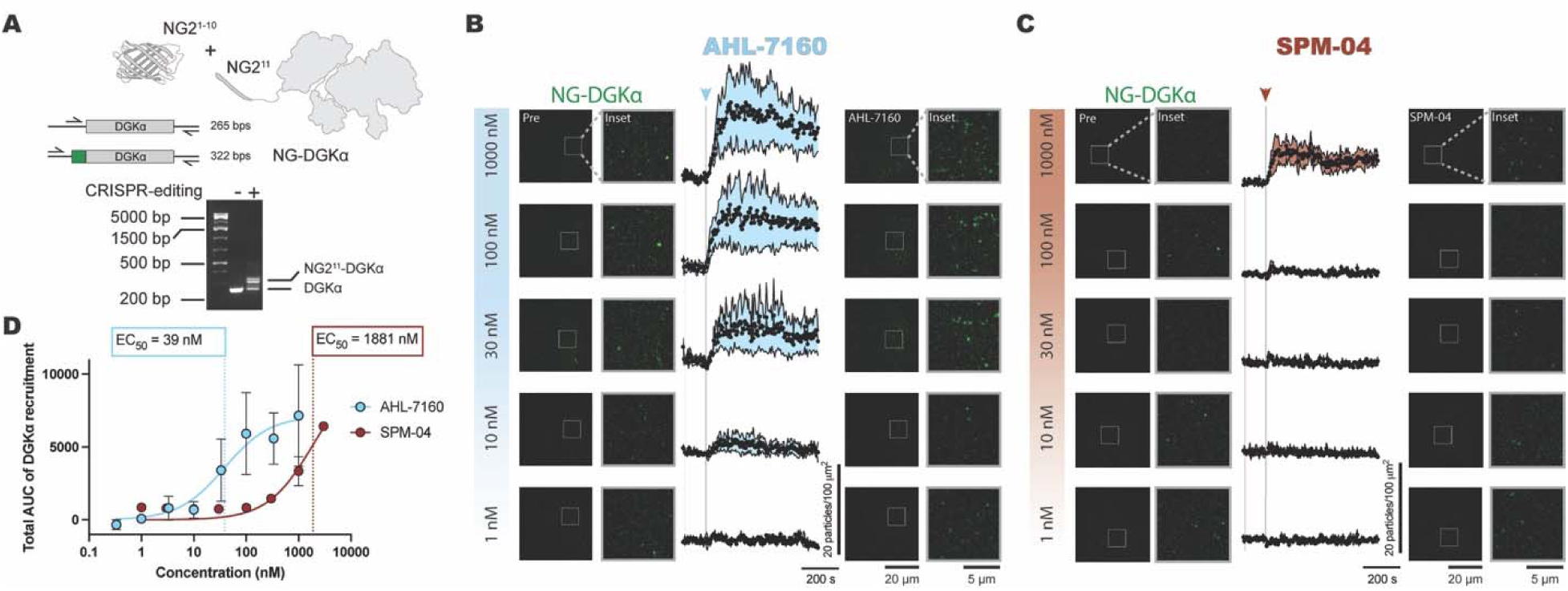
Stereoselective plasma membrane recruitment of endogenous DGKα. (A) PCR was used to confirm knock-in of NG2^11^ into DGKα genomic DNA. Primers bound the genomic DNA on either end of the Cas9 cut site. Placement of the primers and the expected band sizes are as depicted. All knock-ins were confirmed by sequencing the expected bands. (B) Knock-in fluorescent DGKα cells were treated with increasing concentrations of AHL-7160 at 120 s (shown by the arrow/line) and imaged using TIRF microscopy for 720 s. The number of DGKα particles localized in a representative 100 um^2^ area of the plasma membrane were quantified. “Pre” images were taken at –120 s before stimulation and the stimulated images were taken at 600 s. Insets were created for each image as shown. (C) Cells were treated as in B, but using the enantiomer SPM-04. (D) The total areas under the curve of the DGKα responses in B and C were calculated (shown as symbols) and plotted as a dose-response curve. A non-linear fit was created by constraining the bottom value to 0 and EC_50_ > 0 (shown as lines). AHL-7160 df = 21, R^2^ = 0.4875, 95% CI of EC_50_ is 6 to 226 nM. SPM-04 df = 21, R^2^ = 0.8173, 95% CI of EC_50_ is 719 to 7248 nM. Data plotted is the mean of means with SEM of 2-3 independent experiments with a total of 9-17 cells.

Next, we utilized this translocation assay to determine isozyme selectivity of chiral ligands against endogenous DGKs in cells. We CRISPR-edited either a NG tag or a StayGold (SG) tag^41^ to at least one representative member of each DGK subtype and confirmed knock-in by PCR and sequencing (Figure S4). For isoforms where PCR was not efficient, we confirmed specificity of the fluorescence seen with confocal imaging after siRNA-mediated knockdown of endogenous DGKζ and DGKθ, which largely eliminated detection (Figure S4B and C). As expected, the type 2 (DGKδ-NG, DGKη-NG), 4 (SG-DGKζ) and 5 DGKs (DGKθ-SG) were principally found in the cytoplasm while the type 3 DGK (DGKε-NG) showed endoplasmic reticulum (ER) localization (Figure S5). Treatment with either AHL-7160 or SPM-04 did not affect plasma membrane recruitment of any of the additional DGKs tested, supporting isozyme-selectivity of AHL-7160 activity against native DGKs in cells (Figure 3).

**Figure 3.**
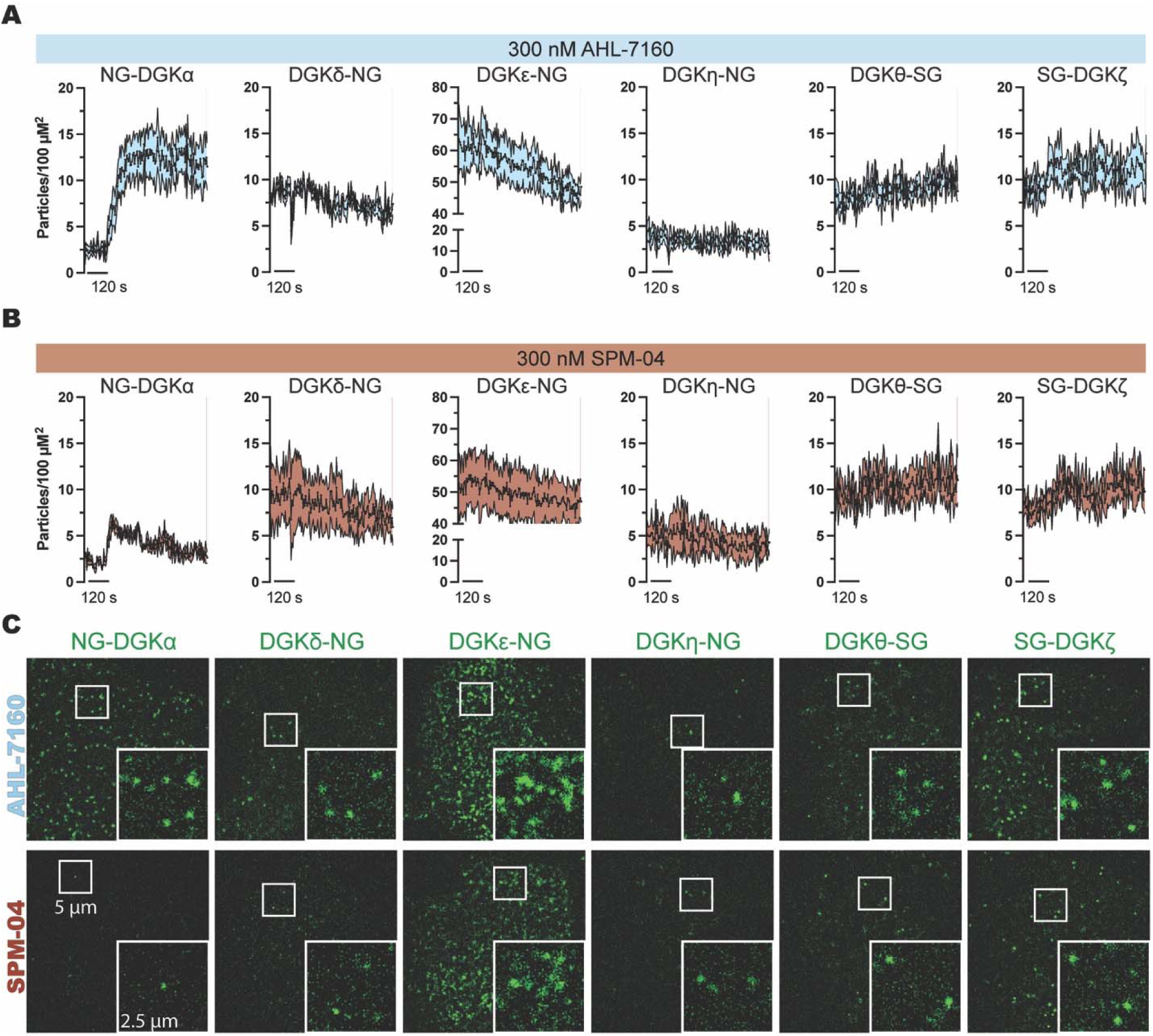
Membrane translocation activity of AHL-7160 is DGK isozyme selective. (A) The DGK knock-in lines were treated with 300 nM AHL-7160 at 120 s as indicated by the scale bar. The number of DGK particles that were recruited to the PM were counted using TIRF microscopy over 720 s. Graphs show the mean of means ± SEM. N = 3-4 experiments with 11-14 total cells. (B) The DGK knock-in lines were treated with 300 nM of SPM-04 enantiomer at 120 s as indicated by the scale bar. The number of DGK particles that were recruited to the PM were counted using TIRF microscopy over 720 s. Graphs show the mean of means ± SEM. N = 3-6 experiments with 12-20 total cells. (C) Representative TIRF images of the DGKs’ responses to SPM-04 or AHL-7160 after 720 s. Insets show individual DGK particles.

### AHL-7160 functions as a molecular glue for stereoselective degradation of DGKα

Next, we utilized the quantitative LC-MS/MS chemoproteomic method, tandem mass tag (TMT)-SuTEx^42^ to determine if AHL-7160 covalently binds DGKα in Jurkat T cells. These studies are important since target engagement between native DGKs and naphthyridinone– and pyridopyrimidinone-based inhibitors in cells has not been demonstrated. In addition, the site of binding for these optimized inhibitors, while approximated using base editing mutagenesis, remains to be fully defined^36^.

Previous studies identified TH211 as a suitable SuTEx probe for quantifying functional sites on endogenous DGKα in Jurkat cells under *in situ* labeling conditions^43^. To test whether AHL-7160 covalently engages DGKα in T cells, we treated cells with this ligand or SPM-04 enantiomer followed by TH211 *in situ* labeling. Cells were harvested, lysed and proteomes subjected to TMT-SuTEx as previously reported^42^ and described in Methods. Using this in-cell competitive activity-based protein profiling (ABPP) approach, we discovered treatment with AHL-7160 (5 µM, 2 hrs) resulted in stereoselective engagement of Y669 located in the DAGKa region of the catalytic domain (competition ratio or CR of 0.4 vs 1 for AHL-7160 compared to SPM-04, respectively; Figure 4A and S6A-B). The K411 site in the DAGKc catalytic region was also stereoselectively engaged by AHL-7160 (CR = 0.6). The detected liganding events were not due to changes in DGKα protein expression (abundance ratios of ∼1, Figure 4B). All liganded sites quantified in TMT-SuTEx studies can be found in Table S1.

**Figure 4.**
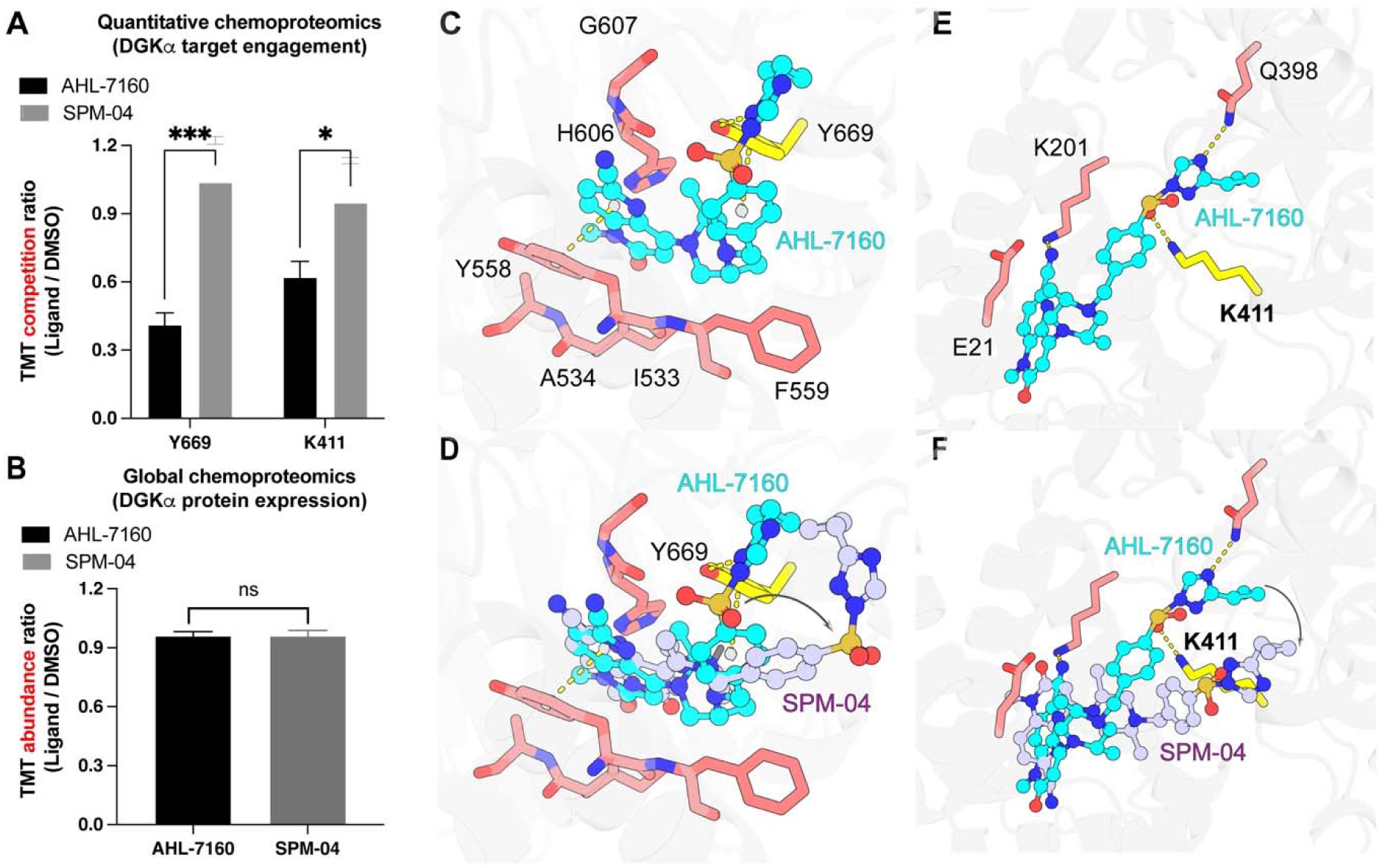
AHL-7160 stereoselectively engages DGKα Y669 and K411 in treated T cells. (A) Competitive TMT-ABPP analysis of proteomes from Jurkat cells pretreated with AHL-7160 or SPM-04 (5 µM, 2 hrs) followed by TH-211 probe labeling (50 µM, 2 hrs) to demonstrate ligand engagement of DGKα Y669 and K411 site. Data shown are representative of n = 3 biological replicates. See Table S1 for the complete proteomic dataset. (B) DGKα protein expression with AHL-7160 and SPM-04 treatment was not affected as measured by global proteomics. (C) Docking of AHL-7160 on the predicted DGKα structure (AF-P23743-F1). Residue Y669 can form a hydrogen bond with the triazole moiety, positioned adjacent to the SuTEx warhead. (D) Overlap of AHL-7160– and SPM-04-DGKα docked structures revealed a conformational change in SPM-04, which positions the SuTEx electrophile further away from the Y669 nucleophilic site. (E) K411 can form a hydrogen bond with the sulfonyl group, positioned adjacent to the SuTEx electrophile in the AHL-7160-DGKα docked structure. (F) Overlap of AHL-7160 and SPM-04 docked at DGKα K411 revealed that SPM-04 adopts an altered conformation, resulting in less favorable accommodation in the binding pocket. Hydrogen bonds are highlighted as yellow dashed lines, and π–π interactions are shown in light pink. The figure was generated using PyMOL 3.1

To investigate the binding mode of AHL-7160, we performed a docking study in the binding pockets on DGKα containing the Y669 and K411 sites. The Y669 site lies within a well-defined pocket while K411 is positioned in a flexible loop domain on the AlphaFold structure of DGKα (AF-P23743-F1, Figure S6A). The docking results showed that AHL-7160 and its enantiomer SPM-04 adopt different poses toward the solvent-exposed pocket. AHL-7160 positions the sulfonyl-triazole group near the Y669 site while SPM-04 places the electrophile further away, providing an explanation for the observed stereoselectivity (Figure 4C, D). Overlap of AHL-7160 and SPM-04 docked at DGKα K411 revealed that SPM-04 adopts an altered conformation, resulting in a less favorable accommodation in the binding pocket (Figure 4E, F).

Considering reversible DGK inhibitors have reported molecular glue activity^36,37^, we tested whether longer exposure to AHL-7160 could mediate DGK degradation in T cells. Extended treatments with AHL-7160 in Jurkat cells resulted in concentration-dependent (0.001-10 µM) degradation of DGKα that could be reversed with the proteasome inhibitor MG-132 (6 hrs, Figure 5A). AHL-7160 degradative activity was highly stereoselective and isozyme-specific as evidenced by inactivity of SPM-04 and its negligible effects on DGKζ protein expression, respectively (Figure 5A). Treatment with the reversible analogs of AHL-7160 did not induce DGKα degradation in Jurkat cells (Figure S6C).

**Figure 5.**
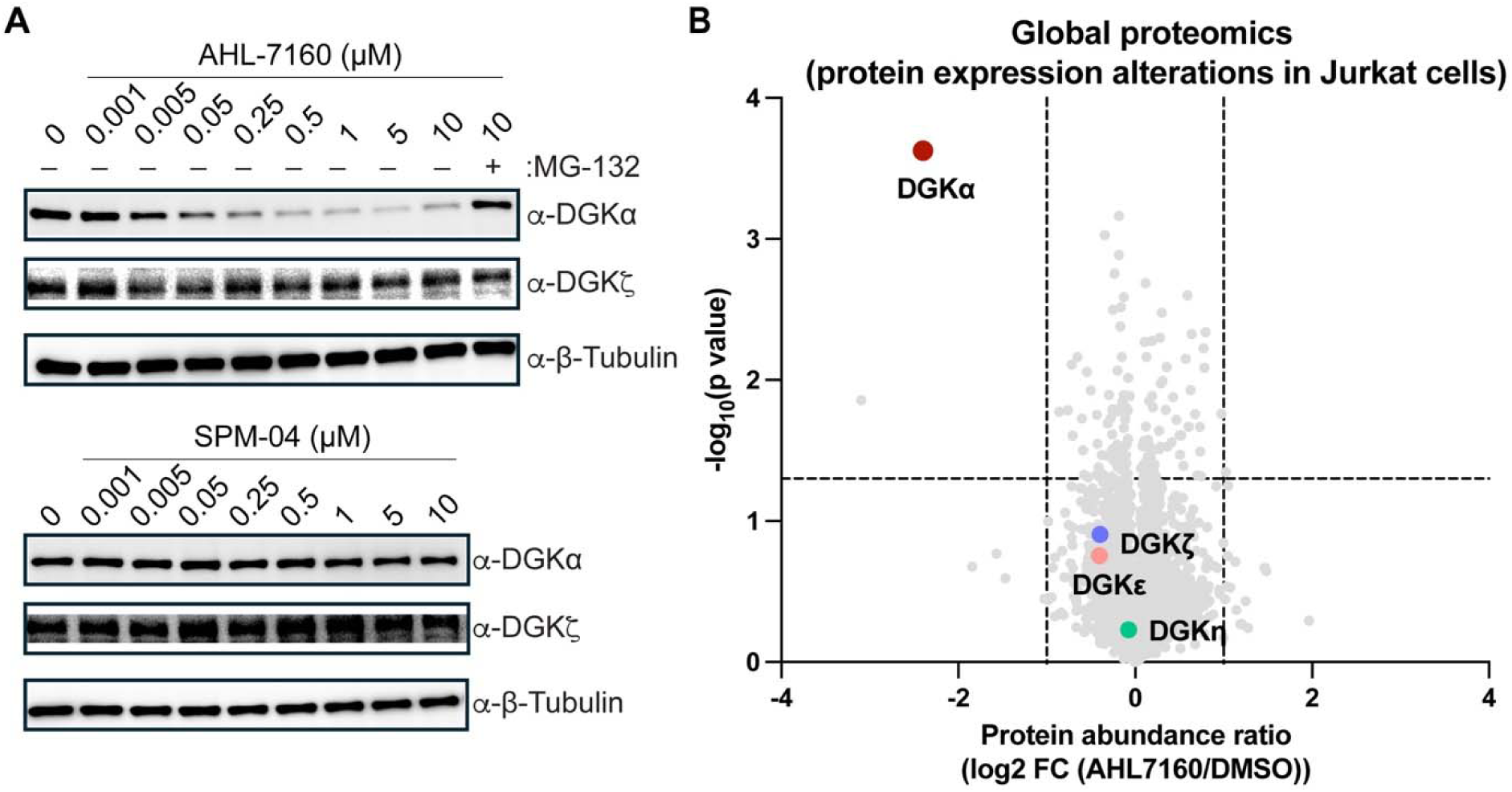
AHL-7160 functions a stereoselective molecular glue degrader of DGKα upon extended treatments. (A) Dose-dependent degradation of DGKα but not DGKζ in AHL-7160-treated Jurkat cells (6 hrs treatment). The molecular glue activity of AHL-7160 in Jurkat cells is stereoselective as evidenced by lack of degradation activity with SPM-04 treatment. (B) Volcano plot showing proteome-wide selective degradation of DGKα by AHL-7160 (5 µM compound, 6 hrs) in Jurkat cells as measured by global proteomics (DIA). A single additional protein (FMC1) was identified as exhibiting significantly downregulated expression in these studies. Data shown are representative of n = 3 biological replicates. See Table S2 for the complete proteomic dataset.

Next, we performed global proteomics to evaluate protein expression changes with extended AHL-7160 treatment in Jurkat cells (5 µM compound, 6 hrs). Remarkably, we identified DGKα as the most significantly downregulated protein (∼80%) with only a single additional protein (FMC1) showing decreased abundance across ∼6,800 total proteins quantified (DGKα abundance ratio = 0.19 from AHL-7160/DMSO treatment, *p* <0.05; Figure 5B). We quantified protein expression of several DGK family members including DGKε, DGKη, and DGKζ and did not observe any changes from AHL-7160 treatment. Importantly, protein expression of ∼280 kinases detected in these global proteomic studies was not affected by AHL-7160 treatment (Table S2).

In summary, we identified AHL-7160 as a site-directed, stereoselective molecular glue degrader of DGKα that exhibited proteome-wide selectivity under extended treatment times in Jurkat T cells.

### AHL-7160 stereoselectively blocks phosphatidic acid production in cells

DGKs convert cellular DAG to PA with preliminary evidence for lipid substrate preference^39,44^. These data, generated primarily through LC-MS lipidomics analyses, provide valuable insights into lipid profiles, while also highlighting the opportunity for complementary analyses that capture dynamic lipid regulation in living cells. Here, we assessed the effects of covalent DGKα chiral ligand treatment on PA production and dynamics in cells using a genetically encoded lipid biosensor.

To monitor direct DAG to PA conversion by DGKα in cells, we established a chemically inducible system to recruit FKBP-PI-PLC and FKBP-DGKα to mitochondrial membranes via rapamycin-mediated dimerization with FRB-Fis1 tail^45^ (Figure 6A). A PA biosensor (PILS-Nir1) was used to detect mitochondrially produced PA using PILS-Nir1 intensity at mitochondria compared to the whole cell (i.e., PILS-Nir1 mito/cell intensity ratio). Addition of rapamycin resulted in a rapid increase in mitochondrial PA as determined by the change in PA biosensor intensity (increase from 1 to 3 PILS-Nir1 mito/cell intensity ratio). Co-treatment of rapamycin with AHL-7160 inhibited PA biosensor intensity at mitochondria, and the potency of this blockade was substantially reduced with SPM-04 (IC_50_ = 340 vs 3200 nM, respectively; Figure 6B-D, Figure S7). We also used the PA biosensor to monitor PA produced at the plasma membrane in response to carbachol (CCh) stimulation and found PILS-Nir1 translocation to the plasma membrane was inhibited with AHL-7160 but not the less active control SMS-55 (Figure S8).

**Figure 6.**
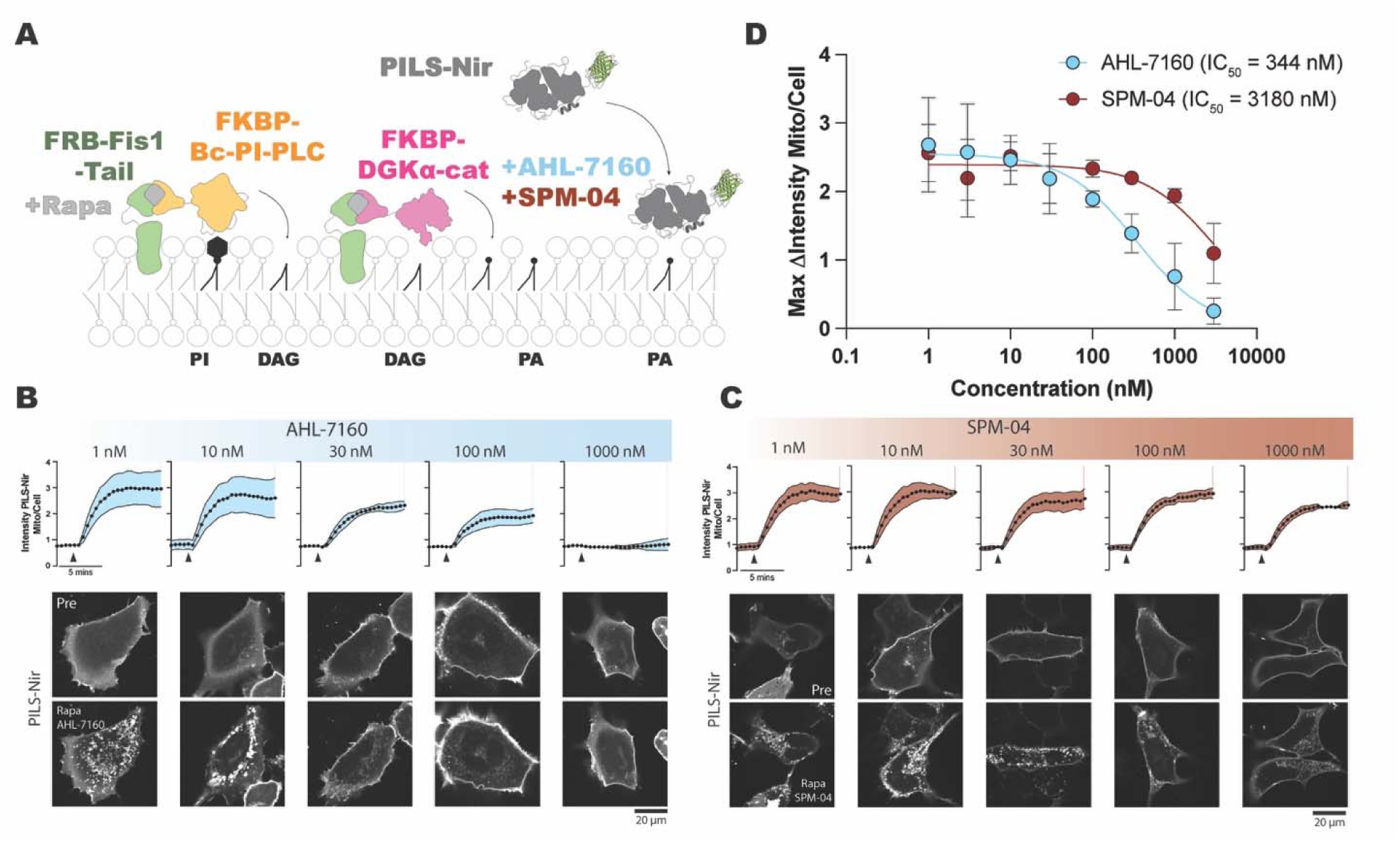
DGKα covalent ligand treatment stereoselectively blocks cellular phosphatidic acid production. (A) Schematic of the chemically inducible system used to make PA at the mitochondria which can be monitored with the PA biosensor PILS-Nir1. (B) HEK293A cells expressing the chemically inducible dimerization system were treated with the concentrations of AHL-7160 shown plus a constant dose of rapamycin. Cells were stimulated at 2 min (denoted by the arrow). The ratio of PILS-Nir1 intensity at the mitochondria to the intensity throughout the whole cell was quantified as a readout of PA levels produced. “Pre” images were taken at –2 min before stimulation and the stimulated images were taken at 10 min. (C) Cells were treated as in B, but using SPM-04. (D) The maximum change in PILS-Nir1 mito/cell intensity ratio was calculated and plotted as a dose-response curve (shown as symbols). A non-linear fit was created by constraining the bottom value to 0 and IC_50_ > 0 (shown as lines). SPM-04 df = 21, R^2^ = 0.3668, IC_50_ 95% CI is 1376 to 10805 nM. AHL-7160 df = 22, R^2^ = 0.6106, IC_50_ 95% CI is 120 to 1048 nM. Data plotted is the mean of means with SEM of 2-3 independent experiments with a total of 25-42 cells.

Collectively, our findings demonstrate AHL-7160 treatment results in stereoselective blockade of DGKα-mediated PA production in different subcellular regions of treated cells.

### AHL-7160 potentiates activation and cytokine response of T cells

DGKα and DGKζ are intracellular checkpoints of T cell signaling by metabolizing lipid messengers necessary for TCR activation^46,47^ (Figure S1A). DAG-mediated signaling through the mitogen-activated protein kinase (MAPK) pathway is reported to be regulated by DGKα and DGKζ^14,19,20,46^. We used anti-CD3 and –CD28 activation conditions tailored for detection of MAPK signaling potentiation as measured by levels of phosphorylated-ERK1/2 (pERK1/2, see Methods for additional details). Treatment with AHL-7160 resulted in concentration-dependent and stereoselective potentiation of pERK1/2 in Jurkat T cells, (Figure 7A).

**Figure 7.**
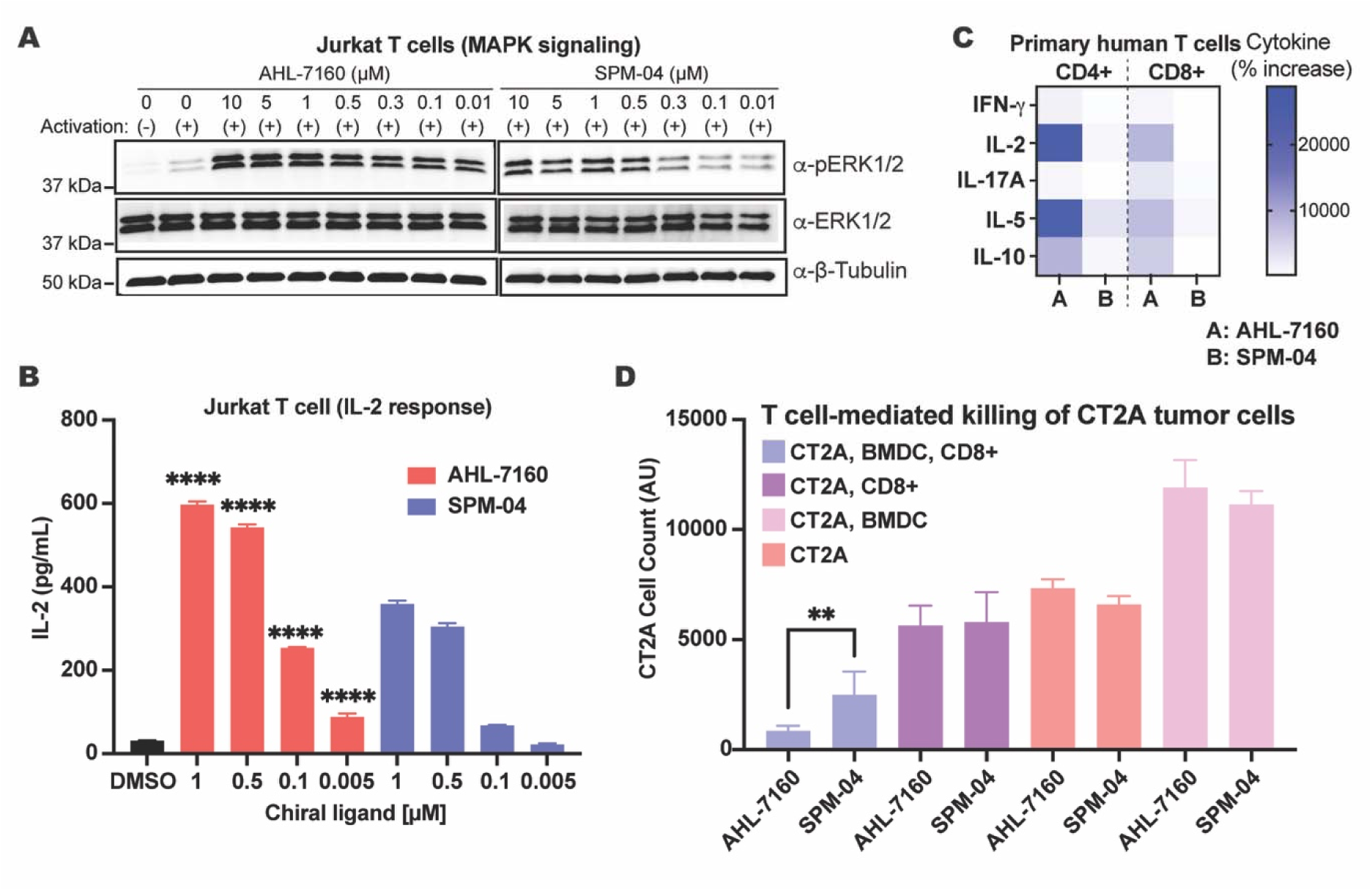
AHL-7160 potentiates T cell activation and killing of tumor cells. (A) Western blot analyses show dose-dependent potentiation of ERK1/2 phosphorylation in AHL-7160-treated Jurkat cells. The reduced activity of SPM-04 supports stereoselective activity of AHL-7160. See Experimental Methods for sub-threshold Jurkat T cell activation conditions using anti-CD3 and –CD28 antibodies. (B) Dose dependent IL-2 production in anti-CD3/CD28-activated Jurkat cells pretreated with AHL-7160 or SPM-04 at the indicated concentrations (30 min). (C) Heat map showing differential cytokine response in primary human CD8+ and CD4+ T cells treated with AHL-7160 or SPM-04 (normalized to DMSO). (D) Co-culture of luciferase-expressing CT2A glioblastoma cells with antigen presenting bone marrow-derived dendritic cells (BMDCs) and CD8+ T cells treated with AHL-7160 or SPM-04. Data shows significant and stereoselective T-cell mediated CT2A killing with AHL-7160 treatment (4 µM compound shown; see Figure S9B for dose-dependent studies).

Next, we tested whether the cytokine response was affected by AHL-7160 treatment in T cells. Akin to pERK1/2, treatment with AHL-7160 resulted in concentration-dependent and stereoselective enhancement of IL-2 production in anti-CD3/CD28-activated Jurkat cells (Figure 6B). We expanded analyses to primary human T cells by testing AHL-7160 activity in CD8+ and CD4+ T cells isolated from peripheral blood mononuclear cells (PBMCs) from healthy donors. T cells were stimulated with anti-CD3 and anti-CD28 in the presence of either AHL-7160 or SPM-04. After 24 hours of stimulation, supernatant was collected and cytokine production was measured by Luminex.

Multiplexed Luminex quantification revealed a coordinated increase in Th1-associated IFN-γ and IL-2, Th2-associated IL-5, Th17-associated IL-17A, and the anti-inflammatory cytokine IL-10, indicating broad immune activation in both CD4+ and CD8+ T cells treated with AHL-7160. Importantly, the enhanced cytokine response in primary T cells was stereoselective as evidenced by reduced or negligible effects with equivalent SPM-04 treatments (Figure 7C). This cytokine signature supports a non-polarized, multi-effector T cell response, rather than selective Th1, Th2, or Th17 skewing elicited by DGKα chiral ligand treatment.

### Covalent DGKα inhibitor treatment activates T cell killing of glioblastoma cells

Next, we asked whether the potentiated T cell activation from AHL-7160 treatment translated into enhanced cytotoxic activity towards tumor cells in an *in vitro* co-culture system. Splenocytes and bone marrow cells were isolated from C57BL/6 mice. Splenic CD8+ T cells and bone marrow-derived dendritic cells (BMDCs) were prepared as described in Supplementary Methods. We evaluated the ability of DGKα chiral ligands to enhance CD8+ T cell-mediated killing of CT2A glioblastoma (GMB) cells (Figure S9A). CT2A is an aggressive and poorly immunogenic mouse GBM cell line that has been used for modeling human GBM resistance to immune checkpoint blockade^48,49^. Thus, CT2A can be used to evaluate compounds for immunotherapy potential.

We tested the effects of DGKα inhibitor (varying concentrations of AHL-7160) on luciferase-expressing CT2A cells (CT2A-luc) and found no significant effects on proliferation compared with DMSO vehicle (Figure 7D). When administered in a co-culture format (CT2A-luc, BMDCs, CD8+ T cells), we observed concentration-dependent killing of CT2A-luc cells in the presence of AHL-7160 (∼70% blockade at 4 µM, Figure 7D). Stereoselectivity of CD8+ T cell-mediated killing of CT2A-luc cells by AHL-7160 was most apparent at lower concentrations when compared to SPM-04 (51% vs 9% for AHL-7160 compared to SPM-04, respectively, at 2 µM treatment; Figure S9B).

## DISCUSSION

Stereoselective recognition is increasingly adopted in chemoproteomic screens to identify ligands with features more suitable for direct cell biological evaluation^50–62^. Ligands that stereoselectively engage cysteines, tyrosines and lysines have been reported, highlighting the potential of this approach. Expanding the diversity of electrophilic scaffolds amenable for chiral ligand development offers further opportunities to expedite ligand discovery. Here, we developed chiral SuTEx ligands for potent and stereoselective degradation of DGKα to block a key lipid metabolic checkpoint and potentiate T cell activation and effector functions.

We embedded the SuTEx electrophile into a high affinity drug scaffold to enable development of a first-in-class covalent DGKα-selective inhibitor (Figure 1). The comparable potency of AHL-7160 with reported reversible DGKα inhibitors^34,36^ was initially surprising given the bulky nature of the SuTEx warhead compared to smaller and more widely adopted electrophilic groups such as acrylamides^63^. Although additional examples are needed to assess generalizability, the current findings support feasibility for introducing the SuTEx electrophile at later stages of targeted covalent ligand development. Importantly, the stereoselectivity of DGKα covalent ligands was key for assessing on-versus off-target activity in complex membrane translocation, cell signaling and T cell co-culture assays.

A covalent ligand format facilitated target engagement studies for accurate site of binding identification on endogenous DGKα in T cells. The ability to perform competitive ABPP studies in living T cells using TH211was important for capturing ligand competition events in native environments. These findings provided evidence in support of covalent binding at the conserved catalytic (DAGKc) and accessory subdomain (DAGKa) of the DGKα catalytic domain (Figure 4). Our findings along with prior reports identifying DAGKa engagement by first-(ritanserin^39^) and second-generation (BMS-986408^36^) reversible DGK inhibitors positions this less conserved region in the catalytic domain as a ligandable hotspot for inhibitor design. Notably, prolonged treatment with AHL-7160 elicited proteasome-dependent DGKα degradation with proteome-wide selectivity (Figure 5). These findings support feasibility for developing highly selective molecular glues that covalently target residues beyond cysteine.

We established a TIRF-based membrane translocation assay using fluorescently tagged endogenous DGKs and demonstrated AHL-7160 stereoselectively induced DGKα translocation in an isozyme specific fashion (Figure 2 and 3). These findings are significant since DGKs, with the exception of membrane-bound DGKε, can reside in the cytosol but must translocate to membranes to catalyze biochemical functions^30^. Our approach addresses the differential expression and activity of DGK isozymes in cells, which makes assessing ligand selectivity across the DGK family particularly challenging^30,31^. Consequently, functional assays using purified DGKs or recombinant DGK-expressing lysates are used as a surrogate approach, which may not reflect endogenous protein behavior^35,36^.

The stereo– and proteome wide-selectivity of AHL-7160 was important for assessing metabolic and signaling activity in cells treated with DGKα chiral ligands. We evaluated the effects of AHL-7160 on DGKα metabolic function using a genetically encoded biosensor that monitors PA dynamics in cells. These studies revealed potent and stereoselective blockade of PA production at different subcellular locations (Figure 6). These alterations in metabolic activity were mirrored by potent and stereoselective potentiation of cytokine response of CD4+ and CD8+ primary human T cells. While additional studies are needed to fully understand the observed multi-effector cytokine response, we were able to correlate this signaling potentiation with enhanced CD8+ T cell-mediated killing of a poorly immunogenic GBM cell line (Figure 7).

The present study opens multiple avenues for future exploration on DGK ligand discovery. Evaluating the ability of AHL-7160 to activate T cell mediated cytotoxicity against a broader tumor cell line panel and *in vivo* will be important to further assess the immunotherapy potential of covalent DGKα ligands. We recognize that AHL-7160 functions as both a DGKα inhibitor and degrader and additional mechanistic studies are warranted to discern the contributions of these respective activities to the underlying pharmacology detected in cells. Whether covalent DGKζ inhibitors could be developed using a similar strategy, i.e., via late stage SuTEx functionalization, represents another opportunity for further exploration. While recent studies have focused on DGKα and DGKζ inhibitors, developing tool compounds for additional DGK isozymes is essential to broaden our understanding of this lipid kinase family beyond oncology^18,30,31^. Finally, although molecular docking provided initial clues to stereoselectivity of DGKα chiral ligands, structural studies of compounds bound to DGKs will facilitate future efforts in structure-enabled ligand design.

In summary, we demonstrate the SuTEx electrophile can facilitate development of stereo– and proteome wide-selective DGKα ligands for targeting a key lipid metabolic checkpoint with applications for immuno-oncology.

## ACKNOWLEDGMENTS

We thank all members of the Hsu lab for helpful discussions and review of the manuscript. This work was supported by the National Institutes of Health grant nos. NS126265 (B.P.), GM144472 (K.-L.H.), DA043571 (K.-L.H.), AI169412 (K.-L.H.), R35GM119412 (G.R.V.H), 1F31HL170755-01 (C.C.W.), T32GM139796 (W.J.W.), University of Virginia Cancer Center (NCI Cancer Center Support Grant No. 5P30CA044579-27 to A.H.L., other to B.P.), University of Texas at Austin startup funds, the Mark Foundation for Cancer Research (Emerging Leader Award to K.-L.H.), a Research Grant Award from The Welch Foundation (F-2143-20230405 to K.-L.H.), and a Recruitment of Rising Stars Award from CPRIT (RR220063 to K.-L.H.).

