## Supplementary Information for "Stereoselective Degradation of Diacylglycerol Kinases Potentiate T cell Activation and Tumor Cell Cytotoxicity"

**CONTENTS**:

1. Supporting Figures
2. Supporting Methods
3. Chemical Synthesis
4. NMR Spectra
5. References

### 1. SUPPORTING FIGURES

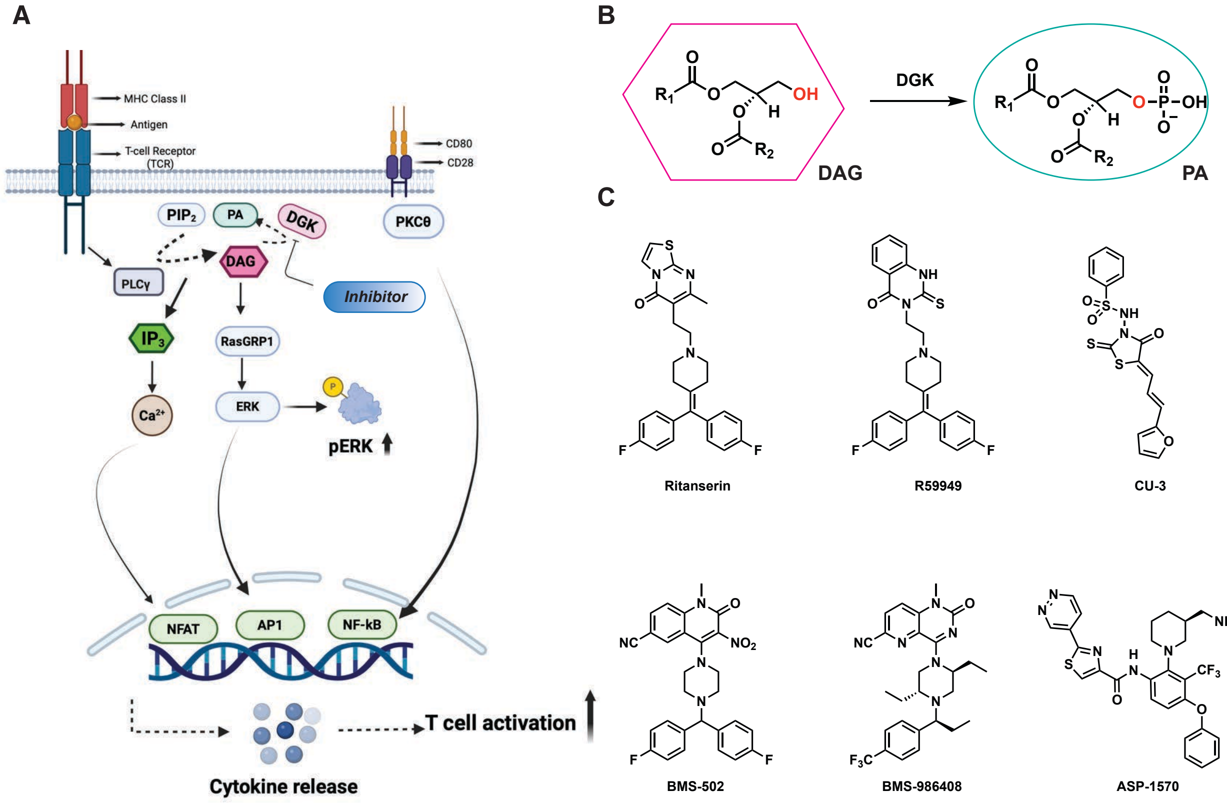


**Figure S1. Diacylglycerol kinases (DGKs) are metabolic checkpoints and pharmacological targets for immunotherapy.** (A, B) T-cell receptor signalling and the role of diacylglycerol kinases (DGKs) in regulating phosphorylation of the key secondary messenger diacylglycerol (DAG) to biosynthesize phosphatidic acid (PA). Pharmacological inhibition of DGKs restores deficient DAG signalling, a feature of the immunosuppressive tumor microenvironment, to enhance T-cell activation, cytokine production, and cytotoxic function. (C) First and second generation DGK inhibitors.

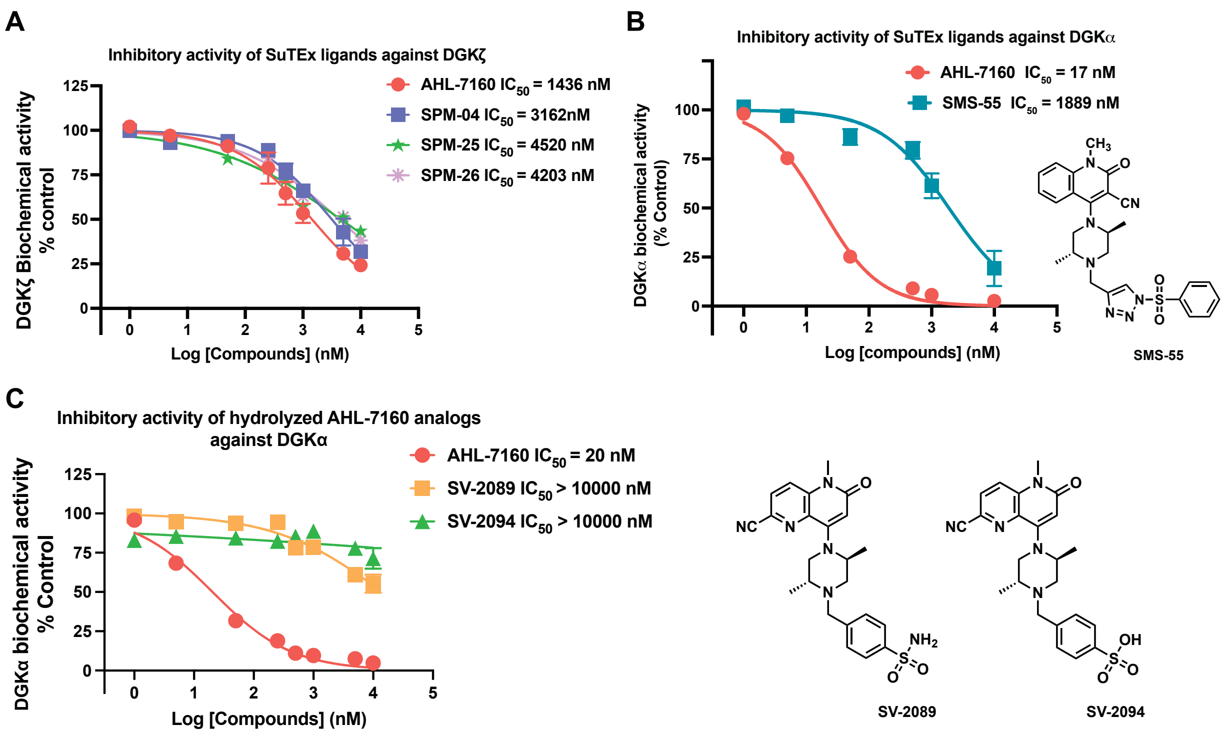


**Figure S2. Activity and covalent binding mechanism of AHL-7160 analogs.** (A) AHL-7160 and its enantiomeric and diastereomeric counterparts namely SPM-04 and SPM-25/SPM-26 show reduced potency against DGKζ as measured by the micelle-based ADP-glo biochemical assay. (B) SMS-55 contains the DGK binding element on the leaving group and shows reduced activity against recombinant DGKα. (C) The reversible analogs of AHL-7160, namely SV-2089 and SV-2094 are largely inactive against recombinant DGKα.

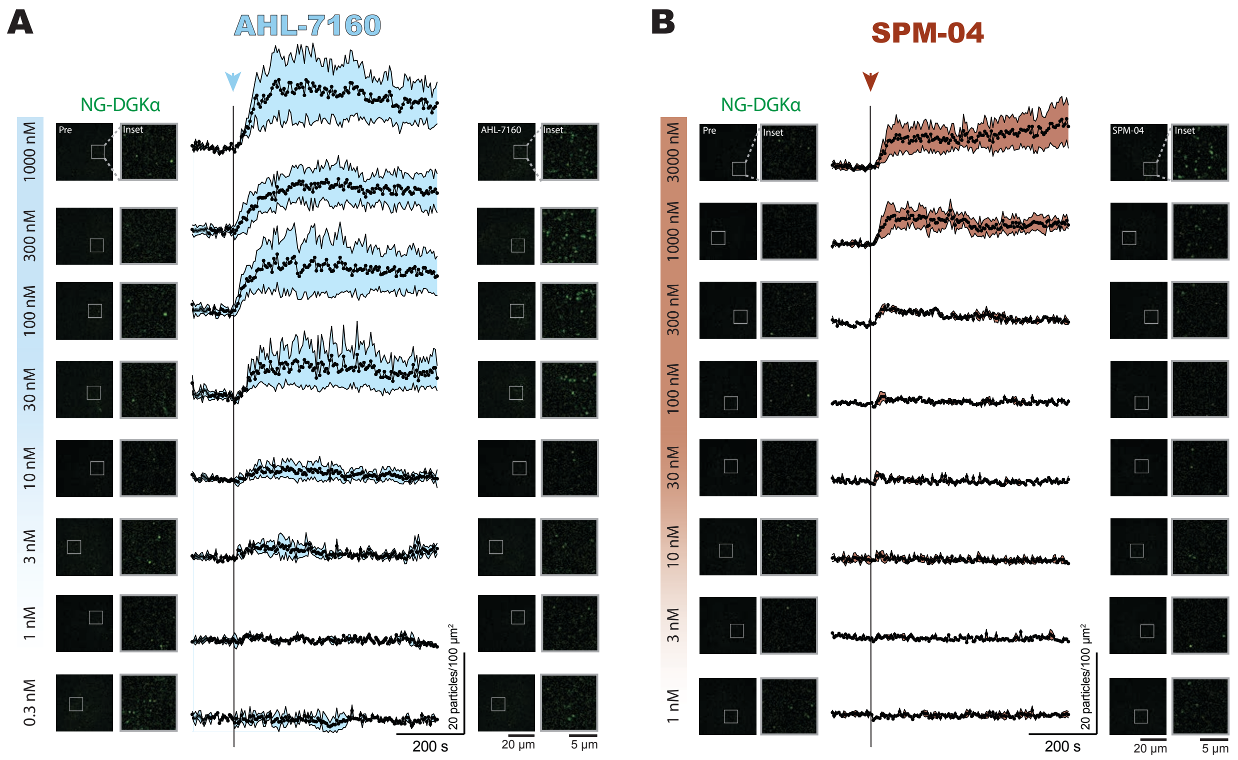


**Figure S3. The full range of doses tested for DGKα membrane recruitment.** Cells expressing NG-tagged endogenous DGKα were treated with the indicated concentrations of AHL-7160 (A) or SPM-04 (B) and DGKα recruitment to the plasma membrane was analyzed as described in Figure 2. This data was used to generate the dose-response curves shown in Figure 2D. The data for the 1 nM, 10 nM, 30 nM, 100 nM, and 1000 nM doses of each drug is replicated from Figure 2.

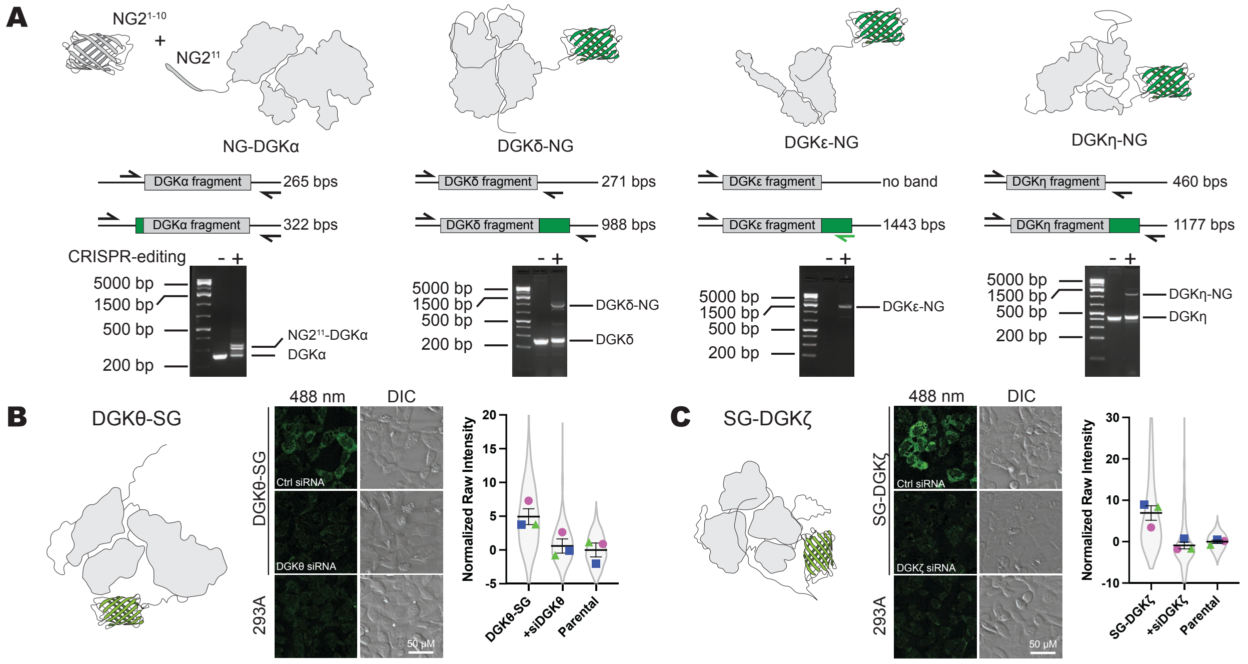


**Figure S4. CRISPR-edited cell lines expressing fluorescent DGK isoforms show AHL-7160 specifically recruits DGKα to the plasma membrane.** (A) PCR was used to confirm knock-in of NG2[11] or full-length NG into DGKα, DGKδ, DGKε, and DGKη genomic DNA. For DGKα, DGKδ, and DGKη, primers bound the genomic DNA on either end of the Cas9 cut site. For DGKε, one primer bound to the NG sequence and the other primer bound to the DGKε genomic DNA. Placement of the primers and the expected band sizes are as depicted. All knock-ins were confirmed by sequencing the expected bands. (B) Knock-in of DGKθ-SG was confirmed using siRNA against DGKθ to show that the fluorescence seen in the 488 nm channel was specific for DGKθ expression. Parental 293A cells were imaged to show baseline autofluorescence, which was set to zero in the graph. Violin plot is the intensity distribution of all cells imaged (DGKθ-SG = 404 cells, siDGKθ = 611 cells, Parental = 666 cells). Symbols show the means of three experimental replicates. Line and error show the grand means with SEM. (C) As in B, but using the SG-DGKζ cell line. Violin plot is the distribution of all cells imaged (SG-DGKζ = 467 cells, siDGKζ = 689 cells, Parental = 609 cells). One experimental replicate of the parental cells was duplicated between B and C. DIC = differential interference contrast.

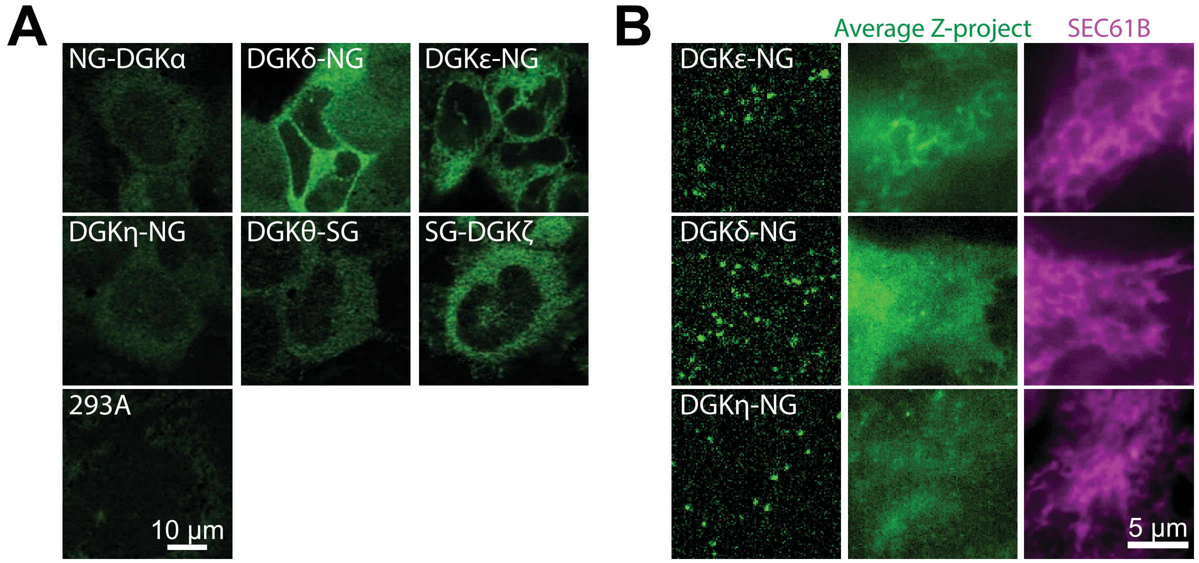


**Figure S5. Basal localization of the CRISPR-tagged DGK isozymes. (A)** Confocal images showing the basal localization of the CRISPR-tagged DGK isozymes. Note the ER-localization of DGKε, while all other isoforms show cytoplasmic localization. Parental 293A cells were imaged to show autofluorescence. (B) The average z-projection of DGKε-NG, DGKδ-NG and DGKη-NG cells shows the ER localization of DGKε-NG compared with the ER marker SEC61B. DGKδ-NG and DGKη-NG are plasma membrane localized negative controls.

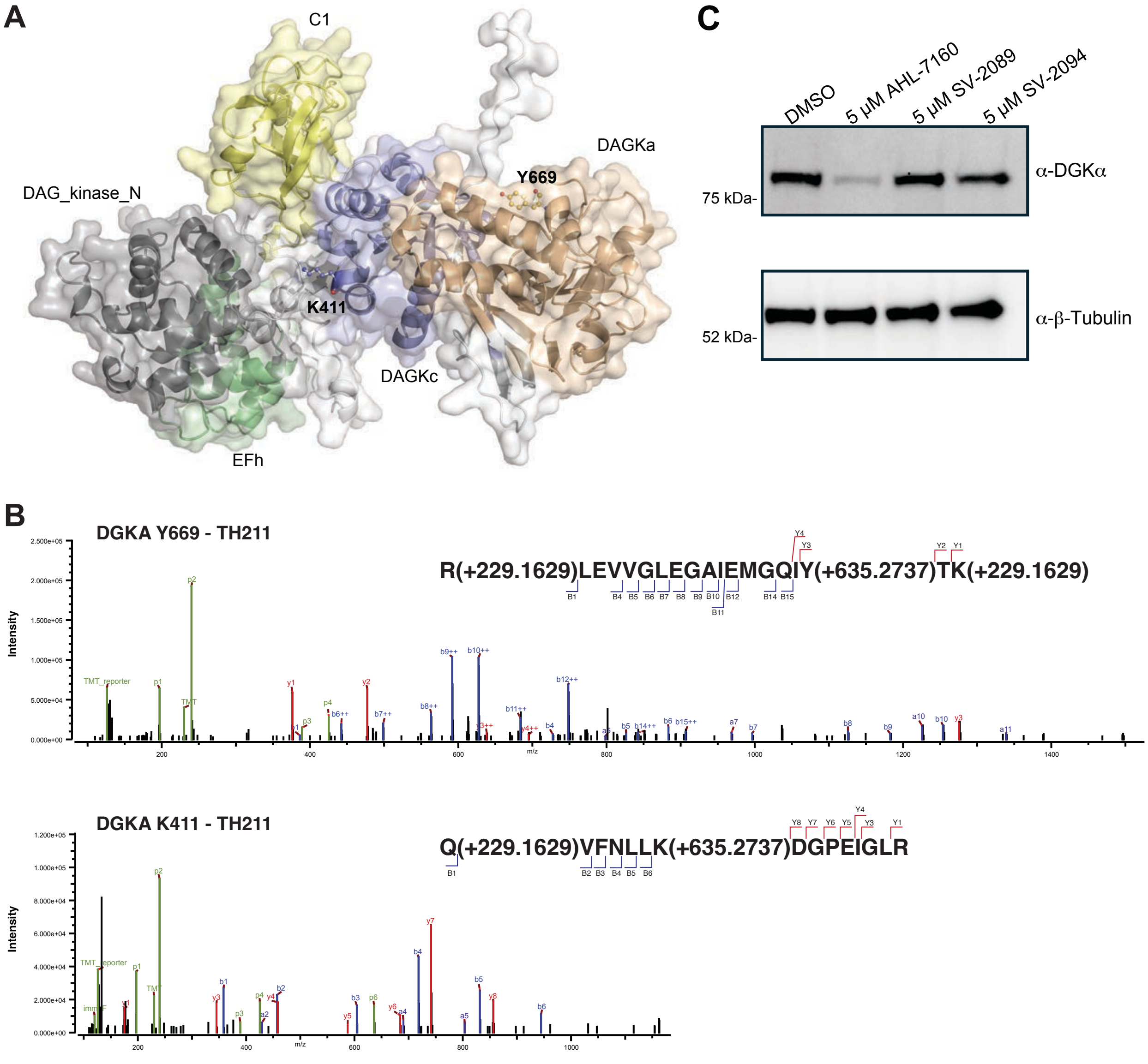


**Figure S6. AHL-7160 binding and docking onto DGKα.** (A) The principal sites of AHL-7160 modification were identified as Y669 and K411, which lies within a well-defined pocket or a flexible loop domain, respectively in the predicted AlphaFold structure of DGKα (AF-P23743-F1). Docking was performed using GOLD 5.1. The figure was generated using PyMOL 3.1. (B) Annotated MS/MS spectra of the TH211 probe-modified peptides corresponding to the Y669 and K411-modified sites. (C) Reversible analogs of AHL-7160 do not induce degradation of endogenous DGKα in T cells.

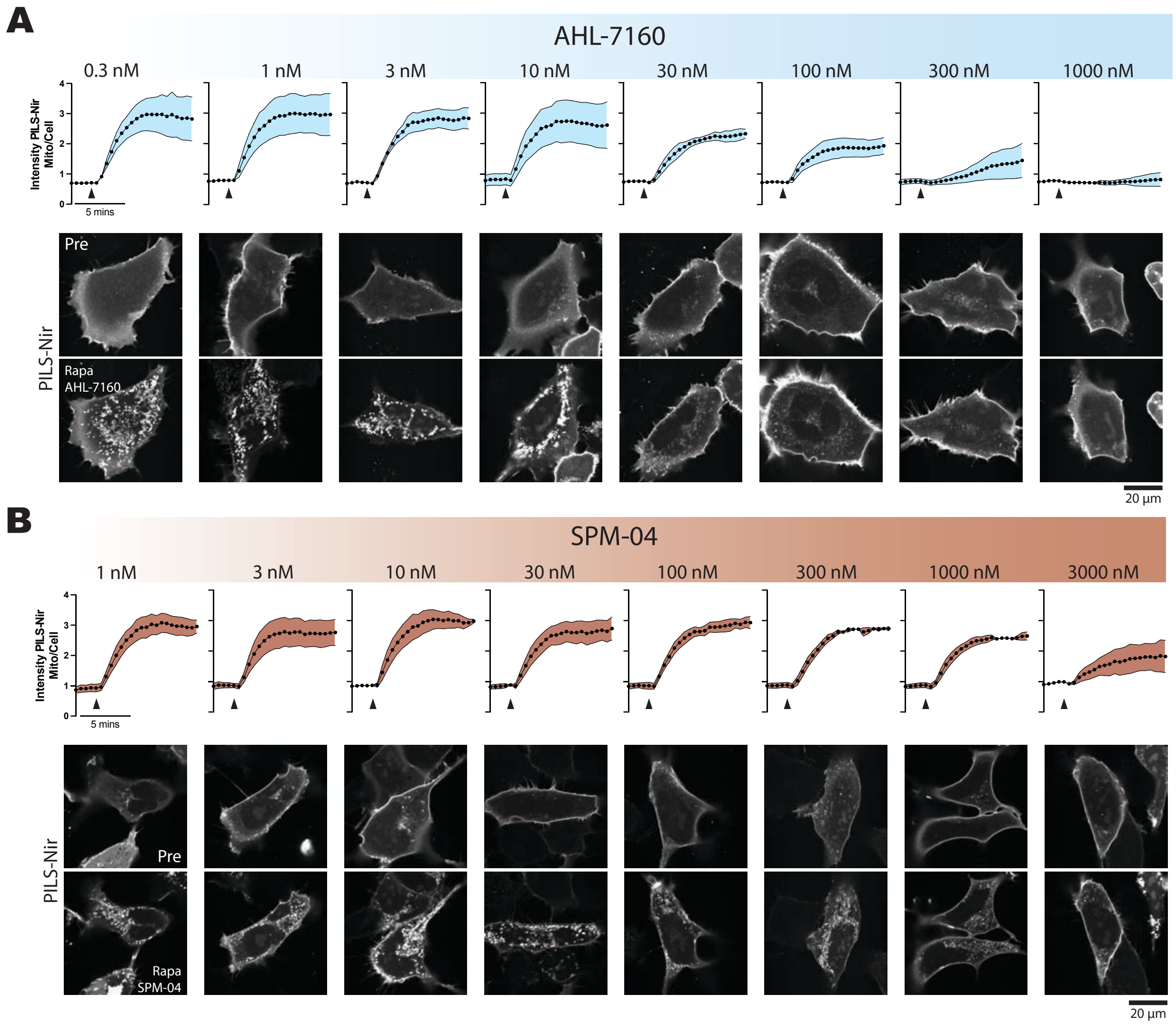


**Figure S7. The full range of doses tested to inhibit FKBP-DGKα.** Cells expressing a chemically inducible dimerization system to make phosphatidic acid at the mitochondria were treated with the given concentrations of AHL-7160 (A) or SPM-04 (B) and PILS-Nir1 recruitment to mitochondria was analyzed as described in Figure 3. This data was used to make the dose-response curves shown in Figure 3C. The data for the 1 nM, 10 nM, 30 nM, 100 nM, and 1000 nM doses of each drug is replicated from Figure 6.

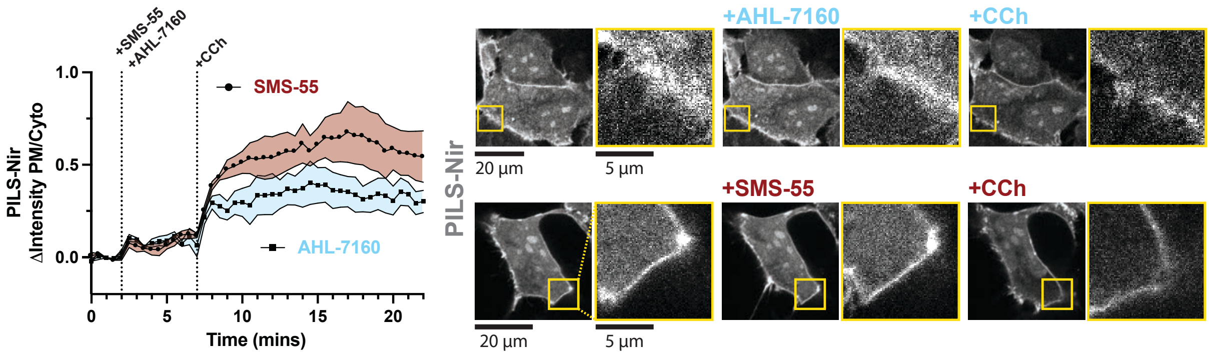


**Figure S8. AHL-7160 inhibits the PILS-Nir response to CCh.** HEK293A cells overexpressing the PILS-Nir biosensor were treated with 300 nM of either AHL-7160 or the negative control compound SMS-55 for 5 min before being treated with 5 mM CCh. The translocation of the biosensor to the plasma membrane (PM) upon phosphatidic acid production was quantified as the change in the fluorescence intensity ratio of sensor at the PM to the intensity within the cytosol. Confocal images from time points 0 mins, 4 mins, and 22 mins are shown with insets highlighting the biosensor at the PM. Graphs show the grand means of 3 experiments with SEM. 31-36 cells were analyzed.

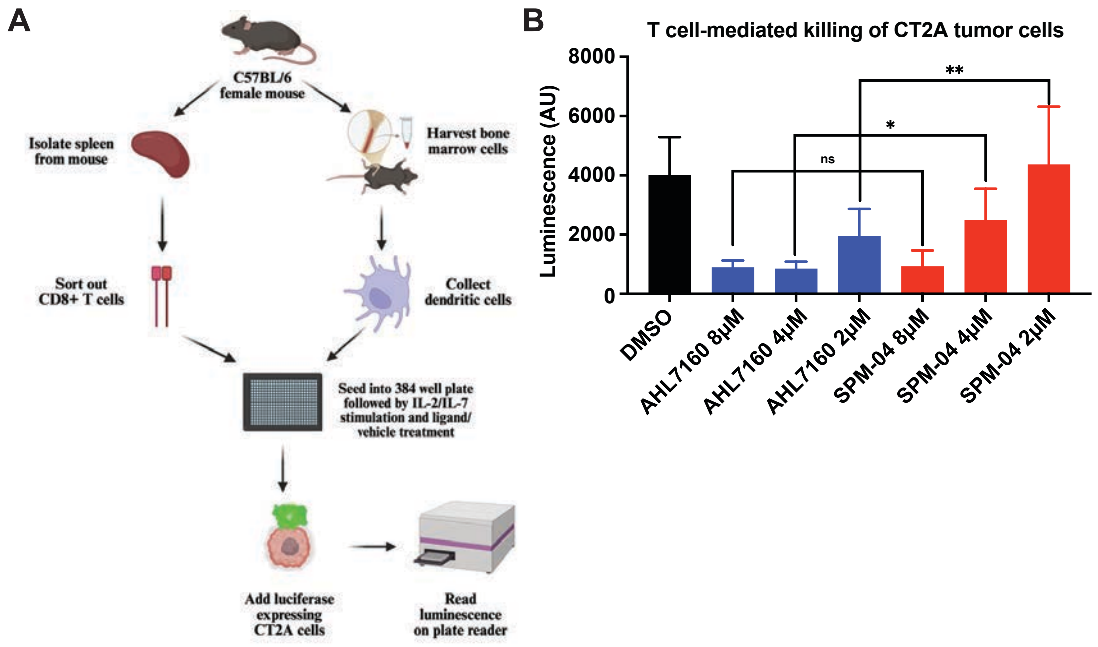


**Figure S9. Co-culture systems for evaluating T cell-mediated killing of CT2A cancer cells.** (A) Experimental scheme for co-culture assay that includes cytotoxic T cells, antigen-presenting cells (bone marrow-derived dendritic cells or BMDCs), and luciferase-expressing CT2A glioblastoma cells. (B) AHL-7160 significantly enhances T-cell mediated CT2A killing at 2 and 4 μM when compared to SPM-04.

### 2. SUPPORTING METHODS

#### Cell culture

HEK293T cells were cultured at 37 °C with 5% $CO_2$ in DMEM supplemented with 10% fetal bovine serum and 1% L-glutamine. Cells were cultured to 80-90% confluency for experimental studies. Jurkat cells were cultured at 37 ºC with 5% $CO_2$ in RPMI supplemented with 10% fetal bovine serum and 1% L-glutamine till cell density reached $1x10^6$ cells/ml for experimental use.

#### Transient transfection

HEK293T cells were used for recombinant protein overexpression following transient transfection procedures as previously described[1]. The wild-type human DGKα and DGKζ plasmid was generated as previously described[2].

#### Cell treatments and lysate preparation

Jurkat cells were treated with DMSO vehicle or compounds in serum-free media for the indicated concentrations and incubation times at 37 °C with 5% $CO_2$ followed by PBS (3 times) washes and cell collection. The cell pellets were lysed by sonication (1 sec pulse, 20% amplitude, 3 times) in PBS in the presence of EDTA-free protease inhibitor cocktail tablet (Pierce). The cell lysates were subject to ultracentrifugation (100,000 x *g*, 45 min at 4 °C) to separate the cytosolic fraction in the supernatant and the insoluble fraction as a pellet. The insoluble pellet was re-suspended in the protease inhibitor-containing PBS by sonication. Protein concentrations in both fractions were measured by the Bio-Rad DC protein assay.

**Western blots**

Western blots were performed as previously described[1]. Anti-FLAG antibody (MilliporeSigma Cat# F1804) was used as the primary antibody for detection of recombinant proteins (1:1000, 5% BSA in 1X TBST) followed by fluorescence detection with a Dylight 550-conjugated secondary antibody (Invitrogen Cat# 84541) using a ChemiDoc imager (Bio-Rad).

**DGK micelle-based biochemical substrate assay**

The ADP-Glo-based DAG phosphorylation substrate assay for measuring DGK biochemical activity was performed as previously described[2]. For jump dilution studies, HEK293T overexpressed hDGKα soluble proteomes were pre-treated with 1 μM AHL-7160 or 25 μM Ritanserin along with DMSO control. After treatment, the reaction mixture was diluted 5X with reaction buffer. The same protocol for the original assay was followed with the amounts of reaction initiator, ADP-Glo reagent and Kinase Detection reagent being scaled up to 5 times to account for the jump dilution. Luminescence was measured on CLARIO Star plate reader.

**Assessing molecular glue activity of DGK ligands**

Jurkat cells were co-treated for 6 hrs with DMSO vehicle, AHL-7160 or SPM-04 at the indicated concentrations in the presence or absence of 10 μM MG-132. Cell lysates were prepared using urea lysis buffer: 50 mM Tris-HCl pH 8.0, 150 mM NaCl, 1 % NP-40, 2M urea and 1 × EDTA-free Protease inhibitor tablet. Changes in DGK protein expression in Jurkat cell lysates (1 mg/mL, 10 μL) were analyzed by western blots as described above. The primary antibodies used for western blots: anti-DGKα antibody (1:6000, ProteinTech cat# 11547), anti-hDGKζ (1:2000, Abcam catalog no. ab239081), and anti-β-tubulin (1:1000, Sigma Aldrich cat#. T5168).

**Quantitation of ERK phosphorylation in activated Jurkat cells**

Jurkat cells were treated with DMSO vehicle or compound at the indicated concentrations and incubation times as indicated in serum free RPMI. Next, cells were activated by anti-CD3 (0.67 nM, Cytek Biosciences Inc cat# 501054552) and -CD28 (3.25 nM, Cytek Biosciences Inc cat# 501054571) antibodies by transferring to antibody coated plates. After treatment, cells were lysed and alterations in total ERK1/2 (Cell Signaling Technology, Inc. 9102S) and phosphorylated ERK1/2 (Thr202/Tyr204, Cell Signaling Technology, Inc. 9101S) were evaluated by western blots. Primary antibodies were diluted 1:1000 in 5% BSA in 1X TBST.

**ELISA to determine IL-2 release in activated Jurkat cells**

Jurkat cells (2.5 x $10^6$ cells/mL) were treated with DMSO vehicle or compound for 30 min followed by activation on anti-CD3 (0.67 nM) and -CD28 (3.25 nM) coated plates for 24 hrs. After treatment, the supernatant was collected and used for ELISA to measure IL-2 secretion following manufacturer protocol (RnD biosciences catalog no. DY20205)

**In vitro human T cell stimulation assay**

96-well round-bottom plates were coated with anti-CD3 antibody (Invitrogen, cat# 16-0037-85) at 0.5 μg/mL in PBS and incubated overnight at 4 °C. $CD4^+$ and $CD8^+$ T cells were isolated from PBMCs using the MojoSort Human CD4 T Cell Isolation Kit (cat# 480009) and the MojoSort Human CD8 T Cell Isolation Kit (cat# 480011), respectively. Before plating, the $CD3^+$ coated wells were washed twice with PBS. A total of 50,000 $CD4^+$ or $CD8^+$ T cells were added per well along with soluble anti-CD28 antibody (Invitrogen, cat# 16.0289.85) at a final concentration of 0.2 μg/mL. Various concentrations of DGKα inhibitors and control reagents were then added to the designated wells. Cells were stimulated for 24 hrs at 37 °C in a humidified 5% $CO_2$ incubator. After 24-hour stimulation, supernatant was collected and stored at -80 °C until the Luminex multi-cytokine analysis.

**Preparation of proteomes for quantitative ABPP site of binding**

Jurkat cells were treated with 5 μM AHL-7160 or SPM-04 for 2 hrs followed by labelling with 50 μM TH-211 probe. Cells were harvested and lysed according to methods previously described[2]. Proteome aliquots (500 μL, 2 mg/mL) were subjected to CuAAC to append Desthiobiotin-azide. Samples extracted, reduced, and alkylated as previously described[3]. Proteins were trypsinized, desalted with Pierce™ Peptide Desalting Spin Columns, TMT labeled, and enriched following previously described methods[3]. After stagetip cleanup, samples were resuspended in 25 μL 0.1% formic acid for LC-MS/MS analysis.

**Preparation of proteomes for global proteomic analysis**

Whole cell lysates from DMSO- or ligand-treated Jurkat cells were normalized to 2.0 mg/mL prior to cleanup by chloroform-methanol extraction as previously reported[2]. Samples were reduced, alkylated, and trypsinized as previously reported[4]. Peptides were TMT labeled, enriched, and cleaned up as previously described[4]. 300 ng peptide aliquots were injected on the LC-MS to quantify global proteomic changes.

**LC-MS/MS data acquisition and analysis for ABPP site of binding**

LC-MS/MS evaluation of TMT-tagged SuTEx modified peptides was performed as previously described on a Vanquish Neo UHPLC coupled to an Orbitrap Eclipse Tribrid mass spectrometer[3]. Spectra were matched and quantified using Proteome Discoverer 3.0 and a PMI Byonic node as previously described[3]. To detect and quantify low abundance probe modified DGKA K411 (855.1337 m/z) and Y669 (800.6861 m/z), we employed parallel reaction monitoring (PRM) in the context of the above-mentioned LC gradient and ms2 acquisition for TMT-tagged SuTEx modified peptides. Spectra matched in the PRM were quantified in Proteome Discoverer 3.0 with filtering for a co-isolation threshold of 80% and Byonic Score cutoff of 100.

**LC-MS/MS data acquisition and analysis for global proteomics**

LC-MS/MS evaluation of TMT-modified peptides was accomplished on a Vanquish Neo UHPLC coupled to an Orbitrap Eclipse Tribrid mass spectrometer as previously described[4]. Spectra matching and protein quantification was performed as previously described[4].

**Data-independent acquisition (DIA) LC-MS/MS analysis of unenriched proteome**

Nano-electrospray ionization–LC–MS/MS analyses were performed using an Orbitrap Exploris 480 mass spectrometer (Thermo Scientific) with the following LC gradient: (A, 0.1% formic acid; B, 0.1% formic acid in 80% ACN): 0-2 min 450 mL/min to 4.5% B, 2-4 min 450 mL/min 20% to B, 4-24 min 300 mL/min to 30% B, 24-74 min 300 mL/min to 45% B, 74-79 min 300 mL/min 50% B, 79-85 min column wash 450 mL/min 99% B. Staggered acquisition using 12 m/z windows, 60 m/z wide from 400-1000 m/z with 120K MS1 resolution and 30K MS2 resolution was used for optimal quantification and identification.

**DIA LC-MS/MS data analysis**

Raw files were converted to .mzML using MSConvert using the Peak Picking filter. Samples were searched in DIA-NN against a modified human protein database (UniProt human protein database, angiotensin I and vasoactive intestinal peptide standards; 40,660 proteins) with the following parameters: up to 2 missed cleavages, up to 2 variable modifications (N-term M excision, C carbamidomethylation, methionine oxidation), peptide length range 7-30 amino acids, precursor charge range 1-4, precursor m/z range 300-1200, automatic inference from first sample for mass accuracy determination, single-pass neural network identification, 1% FDR, and cross-run normalization off (samples were normalized following DIA-NN processing). The protein quantification matrix output from DIA-NN was filtered to exclude proteins with <2 unique peptides contributing to identification, and missingness in >1 (of 3) biological replicates per condition. Protein intensities were log2 transformed, and median (non-zero) normalized prior to Log2 Fold Change (Ligand/DMSO) calculation using the mean normalized intensities. P-values were calculated using a two-sided unpaired T-test between ligand and DMSO intensities. Log2(Fold Change) was plotted against the -Log10(P-value) with a log2(Fold Change) cutoff of 1 and a -Log10(P-value) cutoff of 1.3 ($p < 0.05$).

**Molecular Modeling**

A predicted structure of DGKα (AlphaFold: AF-P23743-F1) was obtained from AlphaFold Protein Structure Database. Residues within 12 Å of the binding site were defined as the docking pocket. Subsequently, compound was docked into this defined pocket using GOLD 5.1 with default parameters and 100 genetic algorithm (GA) runs. For each GA run, a maximum of 125,000 operations were performed. Docking was terminated when the top ten solutions had RMSD values within 1.5 Å. GOLD typically generated 10 poses for each ligand. The top-ranked pose, based on the GoldScore, was selected for display in the figures. The resulting protein–ligand complexes were visualized using the PyMOL Molecular Graphics System (version 3.1).

**In vitro mouse immune cells mediated GBM cell killing assay**

Mouse splenic CD8+ T cells and bone marrow were isolated from C57BL/6 mice as described in Methods. Bone marrow-derived dendritic cells (BMDCs) were differentiated using GM-CSF (20 ng/ml). The immune cells (CD8+ T cells and the BMDCs) were pretreated with DGKα inhibitor AHL-7160 or SPM-04 at 2, 4 and 8 μM for 72 hrs. Then the luciferase expressing mouse GBM cells (CT2A-Luc) were added and incubated at 37 ºC with 5% $CO_2$ for another 72 hrs. The luminescence intensities were measured using GoldBio Luciferase Assay Buffer (TMCA) on a Plate Reader SpectraMax ID3. One-way ANOVA was used for data analyses. All values are mean ± SD.

### 3. CHEMICAL SYNTHESIS

All chemicals used were all reagent grade and used as supplied, except where noted. $^{1}H$ and $^{13}C$ spectra were recorded on a Varian Inova 400 (400 MHz), 600 (600 MHz) spectrometer in $CDCl_3$ with chemical shifts referenced to internal standards ($CDCl_3$: 7.26 ppm $^{1}H$, 77.16 ppm $^{13}C$; $(CD_3)_2SO$: 2.50 ppm $^{1}H$, 40.00 ppm $^{13}C$; $CD_3OD$: 3.31 ppm $^{1}H$, 49.00 ppm $^{13}C$) unless stated otherwise. Splitting patterns are indicated as follows: s, singlet; d, doublet; t, triplet; m, multiplet; br, broad singlet for $^{1}H$-NMR data. NMR chemical shifts ($\delta$) are reported in ppm and coupling constants ($J$) are reported in Hz. High resolution mass spectral (HRMS) data were obtained by an Agilent 6545B LC/Q-TOF (Agilent Technologies, Santa Clara, CA, USA). All reactions were monitored by Thin-Layer Chromatography carried out on precoated Merck silica gel 60 F254 plates (0.25 mm thickness); compounds were visualized by UV light and different stains of a TLC plate. All reactions were carried out under nitrogen or argon atmosphere with dried solvents under anhydrous conditions and yields refer to chromatographically homogenous materials unless otherwise stated. All evaporations were carried out under reduced pressure on Büchi and Heidolph rotary evaporator below 40 °C unless otherwise specified. Compound 1 and compound 4 were synthesized following procedures reported in patents (WO 2020/006018 Al) and (WO 2021/133748 Al).

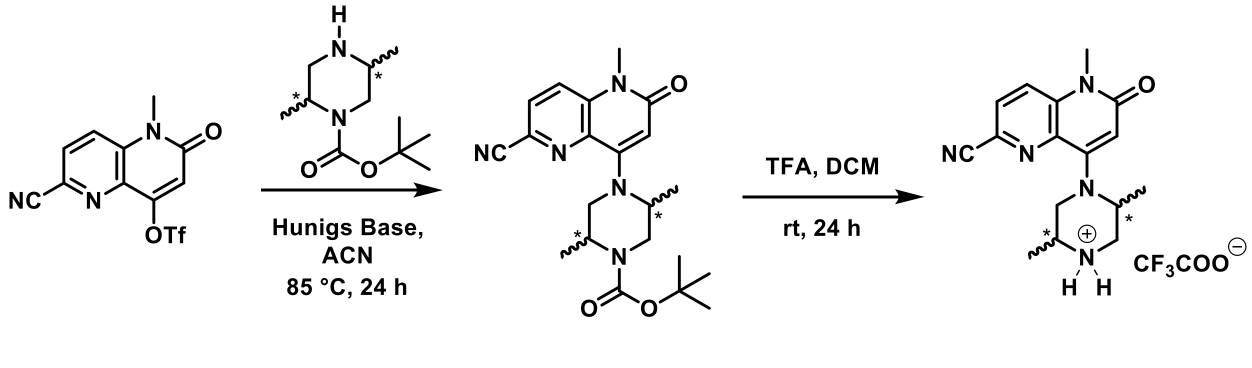


1 2 3

**tert-Butyl(2R,5S)-4-(6-cyano-1-methyl-2-oxo-1,2-dihydro-1,5-naphthyridin-4-yl)-2,5-dimethylpiperazine-1-carboxylate (2a):**

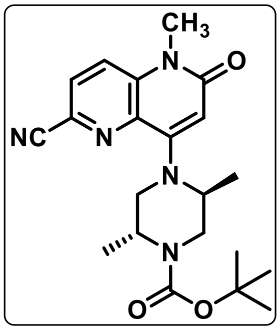

To a stirred solution of **1** (20.36 g, 61.08 mmol) and commercially available *tert*-butyl (2*R*, 5*S*)-2,5-dimethylpiperyzine-1-caroboxylate (13.74 g, 64.13 mmol) was dissolved in anhydrous acetonitrile (400 mL), and Hunig's base (31.9 mL, 183.2 mmol) was added at room temperature. The reaction mixture was stirred at 85 °C for 16 hrs, after which it was cooled down to room temperature and diluted with brine (200 mL) and then extracted with EtOAc (2 x 300 mL). The combined organic layer was dried over $Na_2SO_4$. The residue was purified by Biotage flash chromatography using a gradient of 0% to 85 % EtOAc in *n*-hexane as an eluent to give the desired product **2a** (24.32 g, 92% yield) as a pale yellow solid. **$^1$H NMR** (600 MHz, $CDCl_3$) $\delta$ 7.79 (d, $J$ = 8.7 Hz, 1H), 7.69 (d, $J$ = 8.8 Hz, 1H), 6.12 (s, 1H), 4.60 – 4.29 (m, 2H), 3.89 – 3.63 (m, 2H), 3.62 (s, 3H), 3.49 (d, $J$ = 6.7 Hz, 2H), 1.48 (s, 9H), 1.35 (d, $J$ = 6.8 Hz, 3H), 1.23 – 1.12 (m, 3H).

***tert*-butyl (2*S*,5*R*)-4-(6-cyano-1-methyl-2-oxo-1,2-dihydro-1,5-naphthyridin-4-yl)-2,5-dimethylpiperazine-1-carboxylate (2b):**

To a stirred solution of **1** (1.0 g, 3.00 mmol) and commercially available *tert*-butyl (2*S*, 5*R*)-2,5-

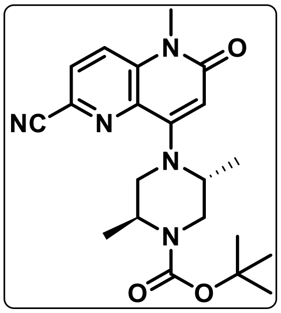

dimethylpiperyzine-1-caroboxylate (0.678 g, 3.15 mmol) was dissolved in anhydrous acetonitrile (20 mL), and Hunig's base (1.6 mL, 9.00 mmol) was added at room temperature. The reaction mixture was stirred at 85 °C for 16 hrs, after which it was cooled down to room temperature and diluted with brine (50 mL) and then extracted with EtOAc (2 x 50 mL. The combined organic layer was dried over $Na_2SO_4$. The residue was purified by Biotage flash chromatography using a gradient of 0% to 85% EtOAc in *n*-hexane as an eluent to give the desired product **2b** (0.750 g, 65 % yield) as a pale yellow solid. $^{1}$**H NMR** (400 MHz, $CDCl_3$) $\delta$ 7.79 (d, $J$ = 8.7 Hz, 1H), 7.69 (d, $J$ = 8.8 Hz, 1H), 6.12 (s, 1H), 4.43 (d, $J$ = 76.9 Hz, 2H), 3.75 (d, $J$ = 16.2 Hz, 2H), 3.62 (s, 3H), 3.58 – 3.40 (m, 2H), 1.48 (s, 9H), 1.35 (d, $J$ = 6.8 Hz, 3H), 1.18 (s, 3H).

***tert*-butyl (2*R*,5*R*)-4-(6-cyano-1-methyl-2-oxo-1,2-dihydro-1,5-naphthyridin-4-yl)-2,5-dimethylpiperazine-1-carboxylate (2c):**

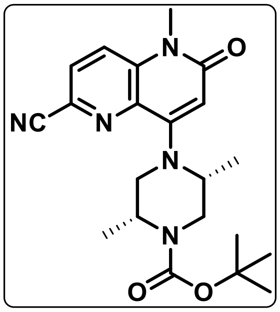

To a stirred solution of **1** (1.0 g, 3.00 mmol) and commercially available *tert*-butyl (2*R*, 5*R*)-2,5-dimethylpiperyzine-1-caroboxylate (0.678 g, 3.15 mmol) was dissolved in anhydrous acetonitrile (20 mL), and Hunig's base (1.6 mL, 9.00 mmol) was added at room temperature. The reaction mixture was stirred at 85 °C for 16 hrs, after which it was cooled down to room temperature and diluted with brine (50 mL) and then extracted with EtOAc (2 x 50 mL). The combined organic layer was dried over $Na_2SO_4$. The residue was purified by Biotage flash chromatography using a gradient of 0% to 85% EtOAc in *n*-hexane as an eluent to give the desired product **2c** (0.791 g, 66.3 % yield) as a pale yellow solid. **$^1$H NMR** (400 MHz, $CDCl_3$) $\delta$ 7.78 (dd, $J$ = 8.7, 0.8 Hz, 1H), 7.67 (dd, $J$ = 8.8, 0.9 Hz, 1H), 6.09 (s, 1H), 4.35 (dt, $J$ = 9.7, 6.0 Hz, 1H), 4.17 (s, 2H), 4.08 (dd, $J$ = 13.9, 6.5 Hz, 1H), 3.61 (s, 3H), 3.19 (d, $J$ = 9.8 Hz, 1H), 2.93 – 2.81 (m, 1H), 1.42 (d, $J$ = 0.8 Hz, 9H), 1.29 (d, $J$ = 5.9 Hz, 3H), 1.24 (d, $J$ = 6.1 Hz, 3H).

***tert*-butyl (2*S*,5*S*)-4-(6-cyano-1-methyl-2-oxo-1,2-dihydro-1,5-naphthyridin-4-yl)-2,5-dimethylpiperazine-1-carboxylate (2d):**

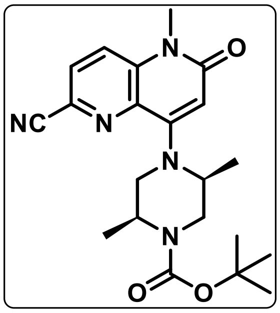


To a stirred solution of **1** (1.0 g, 3.00 mmol) and commercially available *tert*-butyl (2*S*, 5S)-2,5-dimethylpiperyzine-1-caroboxylate (0.678 g, 3.15 mmol) was dissolved in anhydrous acetonitrile (20 mL), and Hunig's base (1.6 mL, 9.00 mmol) was added at room temperature. The reaction mixture was stirred at 85 °C for 16 hrs, after which it was cooled down to room temperature and diluted with brine (50 mL) and then extracted with EtOAc (2 x 50 mL). The combined organic layer was dried over $Na_2SO_4$. The residue was purified by Biotage flash chromatography using a gradient of 0% to 85% EtOAc in *n*-hexane as an eluent to give the desired product **2d** (0.725 g, 60.9% yield) as a pale yellow solid. **$^1$H NMR** (400 MHz, $CDCl_3$) $\delta$ 7.78 (dd, $J$ = 8.7, 0.8 Hz, 1H), 7.67 (dd, $J$ = 8.8, 0.9 Hz, 1H), 6.09 (s, 1H), 4.35 (dt, $J$ = 9.7, 6.0 Hz, 1H), 4.17 (s, 1H), 4.08 (dd, $J$ = 13.9, 6.5 Hz, 2H), 3.61 (s, 3H), 3.19 (d, $J$ = 9.8 Hz, 1H), 2.93 – 2.81 (m, 1H), 1.42 (d, $J$ = 0.8 Hz, 9H), 1.29 (d, $J$ = 5.9 Hz, 3H), 1.24 (d, $J$ = 6.1 Hz, 3H).

**(2*R*,5*S*)-4-(6-cyano-1-methyl-2-oxo-1,2-dihydro-1,5-naphthyridin-4-yl)-2,5-dimethylpiperazin-1-ium-trifluoroacetate (3a):**

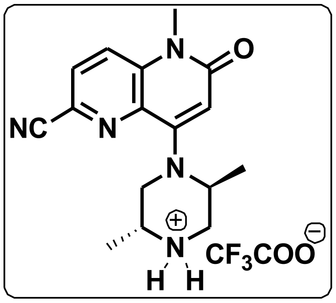


To a stirred solution of **2a** (24.32 g, 64.13 mmol) in DCM (100 mL) and trifluoroacetic acid (51.50 mL, 641.3 mmol) was added dropwise at 0 °C under nitrogen ($N_2$) atmosphere for 20-30 min. The reaction mixture was stirred at room temperature for 16 hrs and then concentrated under reduced pressure. The resultant residue was washed in *n*-Pentane (3 x 150 mL) and $Et_2O$ (1 x 50mL) and then dried under high vacuum to afford the desired final product **3a** (18.5 g, 97% yield) as a white solid. **$^1$H NMR** (400 MHz, $CDCl_3$) *δ* 7.81 – 7.74 (m, 1H), 7.73 – 7.66 (m, 1H), 6.26 (d, *J* = 3.2 Hz, 1H), 3.79 (dd, *J* = 12.0, 3.1 Hz, 1H), 3.69 (s, 1H), 3.63 (d, *J* = 2.7 Hz, 3H), 3.36 – 3.20 (m, 2H), 2.87 – 2.70 (m, 2H), 1.17 (dt, *J* = 6.0, 3.3 Hz, 6H).

**(2*S*,5*R*)-4-(6-cyano-1-methyl-2-oxo-1,2-dihydro-1,5-naphthyridin-4-yl)-2,5-dimethylpiperazin-1-ium-trifluoroacetate (3b):**

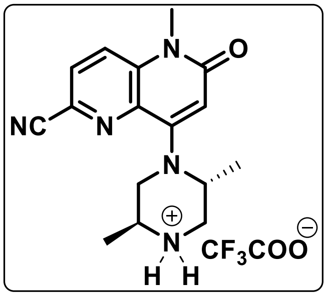


To a stirred solution of **2b** (0.750 g, 1.98 mmol) in DCM (3 mL) and trifluoroacetic acid (1.6 mL, 19.9 mmol) was added dropwise at 0 °C under nitrogen ($N_2$) atmosphere for 20-30 min. The reaction mixture was stirred at room temperature for 16 hrs and then concentrated under reduced pressure. The resultant residue was washed in *n*-Pentane (3 x 10 mL) and $Et_2O$ (1 x 5 mL) and then dried under high vacuum to afford the desired final product **3b** (0.6 g, 92.3% yield) as a white solid. **$^1$H NMR** (400 MHz, $CDCl_3$) *δ* 7.79 (dd, *J* = 8.8, 1.9 Hz, 1H), 7.70 (dd, *J* = 8.8, 1.9 Hz, 1H), 6.28 (d, *J* = 1.9 Hz, 1H), 3.80 (dt, *J* = 12.0, 2.5 Hz, 1H), 3.71 (s, 1H), 3.65 (d, *J* = 1.9 Hz, 3H), 3.32 (t, *J* = 11.9 Hz, 2H), 2.90 – 2.75 (m, 2H), 1.19 (ddd, *J* = 6.0, 4.0, 1.8 Hz, 6H).

**(2*R*,5*R*)-4-(6-cyano-1-methyl-2-oxo-1,2-dihydro-1,5-naphthyridin-4-yl)-2,5-dimethylpiperazin-1-ium-trifluoroacetate (3c):**

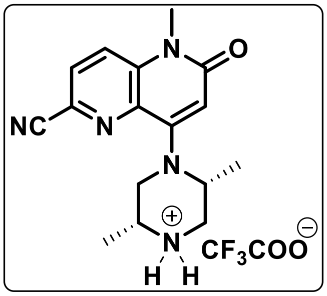


To a stirred solution of **2c** (0.250 g, 0.84 mmol) in DCM (2 mL) and trifluoroacetic acid (1.0 mL, 12.4 mmol) was added dropwise at 0 °C under nitrogen ($N_2$) atmosphere for 20-30 min. The reaction mixture was stirred at room temperature for 16 hrs, and then concentrated under reduced pressure. The resultant residue was washed in *n*-Pentane (3 x 8 mL) and $Et_2O$ (1 x 3 mL) and then dried under high vacuum to afford the desired final product **3c** (0.170 g, 90.9 % yield) as a white solid. **$^1$H NMR** (400 MHz, $CDCl_3$) $\delta$ 7.84 (d, $J$ = 8.8 Hz, 1H), 7.75 (d, $J$ = 8.8 Hz, 1H), 6.20 (s, 1H), 3.75 – 3.66 (m, 1H), 3.64 (s, 3H), 3.55 (d, $J$ = 11.8 Hz, 2H), 3.48 – 3.37 (m, 1H), 3.32 – 3.23 (m, 1H), 2.33 (s, 1H), 1.48 (d, $J$ = 6.4 Hz, 3H), 1.39 (d, $J$ = 7.1 Hz, 3H).

**(2*S*,5*S*)-4-(6-cyano-1-methyl-2-oxo-1,2-dihydro-1,5-naphthyridin-4-yl)-2,5-dimethylpiperazin-1-ium-trifluoroacetate (3d):**

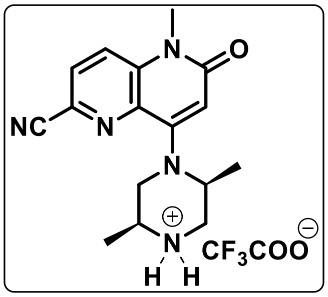


To a stirred solution of **2d** (0.250 g, 0.84 mmol) in DCM (2 mL) and trifluoroacetic acid (1.0 mL, 12.4 mmol) was added dropwise at 0 °C under nitrogen ($N_2$) atmosphere for 20-30 min. The reaction mixture was stirred at room temperature for 16 hrs and then concentrated under reduced pressure. The resultant residue was washed in *n*-Pentane (3 x 8 mL) and $Et_2O$ (1 x 3 mL) and then dried under high vacuum to afford the desired final product **3d** (0.177 g, 94.6 % yield) as a white solid. **$^1$H NMR** (400 MHz, $CDCl_3$) $\delta$ 7.84 (d, $J$ = 8.3 Hz,

1H), 7.75 (d, *J* = 8.4 Hz, 1H), 6.23 (d, *J* = 17.9 Hz, 1H), 3.68 (d, *J* = 9.3 Hz, 1H), 3.65 (s, 3H), 3.59 – 3.47 (m, 2H), 3.43 (q, *J* = 11.5 Hz, 1H), 3.29 (d, *J* = 12.9 Hz, 1H), 2.50 (s, 1H), 1.49 (s, 3H), 1.39 (d, *J* = 7.2 Hz, 3H).

**4-((3-cyclopropyl-1H-1,2,4-triazol-1-yl)sulfonyl)benzaldehyde:**

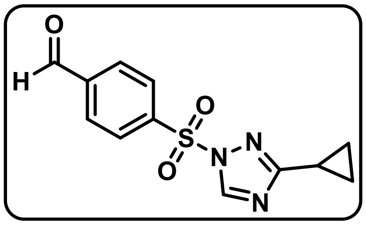


To a stirred solution of 4-formylbenzenesulfonyl chloride (1.00 g, 4.90 mmol) and sodium bicarbonate (823 mg, 9.80 mmol) in anhydrous acetonitrile (10 mL), 3-cyclopropyl-1H-1,2,4-triazole (588 mg, 5.39 mmol) was added. The reaction mixture was stirred at 60 °C for 3 hrs, after which it was cooled down to room temperature. It was diluted with brine (30 mL) and then extracted with EtOAc (2 x 50 mL). The combined organic layer was dried over $Na_2SO_4$. The residue was purified by Biotage flash chromatography using a gradient of 0% to 50 % EtOAc in *n*-hexane as an eluent to give the desired product (0.900 g, 67% yield) as a yellow solid. **$^1$H NMR** (600 MHz, $CDCl_3$) *δ* 10.13 (s, 1H), 8.56 (s, 1H), 8.26 – 8.20 (m, 2H), 8.11 – 8.05 (m, 2H), 2.03 (tt, *J* = 7.8, 5.5 Hz, 1H), 1.00 – 0.97 (m, 4H).

**8-((2*S*,5*R*)-4-(4-((3-cyclopropyl-1H-1,2,4-triazol-1-yl)sulfonyl)benzyl)-2,5-dimethylpiperazin-1-yl)-5-methyl-6-oxo-5,6-dihydro-1,5-naphthyridine-2-carbonitrile (AHL-7160) :**

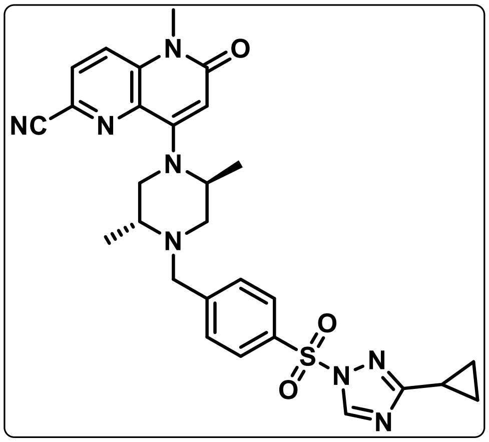


To a solution of **3a** (600 mg, 2.017 mol), in anhydrous DCM (20 mL), 4-(3-cyclopropyl-1H-1,2,4-triazol-1-yl) sulfonyl) benzaldehyde (559 mg, 2.017 mol), sodium cyanoborohydride (380 mg, 6.051 mol) were added. The resulting mixture was stirred at room temperature for 24 hrs, followed by addition of water (50 mL) and then extracted with ethyl acetate (3 × 50 mL). The organic layers were combined and dried over anhydrous $Na_2SO_4$. The residue was purified by Biotage flash chromatography using 95% EtOAc in *n*-hexane as an eluent to give the desired product **AHL-7160** (0.300 g, 28 % yield) as a yellow solid. **$^{1}$H NMR** (600 MHz, $CDCl_3$) $\delta$ 8.55 (d, $J$ = 0.9 Hz, 1H), 8.01 (d, $J$ = 8.0 Hz, 2H), 7.78 (d, $J$ = 8.7 Hz, 1H), 7.69 (d, $J$ = 8.8 Hz, 1H), 7.64 (d, $J$ = 8.0 Hz, 2H), 6.15 (s, 1H), 4.39 (s, 1H), 3.71 (s, 2H), 3.66 (t, $J$ = 11.0 Hz, 2H), 3.63 (d, $J$ = 1.2 Hz, 3H), 3.16 – 3.00 (m, 2H), 2.27 (d, $J$ = 11.6 Hz, 1H), 2.09 – 2.00 (m, 1H), 1.27 (d, $J$ = 6.6 Hz, 3H), 1.21 (d, $J$ = 6.5 Hz, 3H), 0.99 – 0.95 (m, 4H). **$^{13}$C NMR** (100 MHz, $CDCl_3$) $\delta$ 169.76, 162.29, 154.53, 148.25, 145.03, 138.63, 136.52, 134.72, 129.62, 128.72, 128.29, 128.00, 124.42, 122.51, 117.27, 106.91, 58.21, 53.72, 52.41, 51.07, 38.08, 29.15, 14.35, 9.75, 9.16, 8.43. **HRMS** (ESI-TOF) m/z: $[M + Na]^+$ calculated for Chemical Formula: $C_{28}H_{30}N_8O_3SNa$, 581.2054, Found, 581.2061.

**8-((2*R*,5*S*)-4-(4-((3-cyclopropyl-1H-1,2,4-triazol-1-yl)sulfonyl)benzyl)-2,5-dimethylpiperazin-1-yl)-5-methyl-6-oxo-5,6-dihydro-1,5-naphthyridine-2-carbonitrile**

**(SPM-04) :**

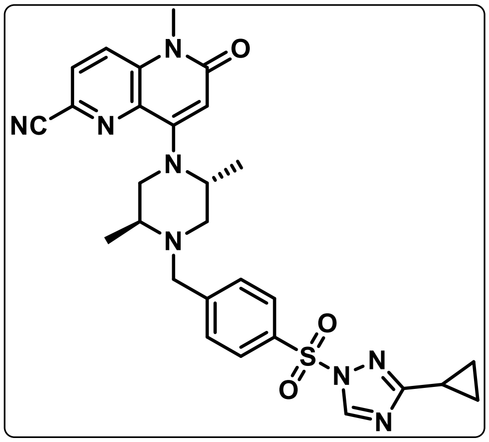

To a solution of **3b (**500 mg, 1.68 mmol), in anhydrous DCM (15 mL)**,** 4-((3-cyclopropyl-1H-1,2,4-triazol-1-yl) sulfonyl) benzaldehyde (466 mg, 1.68 mmol), sodium cyanoborohydride (317 mg, 5.01 mmol) were added. The resulting mixture was stirred at room temperature for 24 hrs, followed by addition of water (50 mL) and then extracted with ethyl acetate (3 × 50 mL). The organic layers were combined and dried over anhydrous $Na_2SO_4$. The residue was purified by Biotage flash chromatography using 95% EtOAc in *n*-hexane as an eluent to give the desired product **SPM-04** (0.186 g, 20 % yield) as a yellow solid. **$^1$H NMR** (400 MHz, $CDCl_3$) $\delta$ 8.55 (s, 1H), 8.02 (d, *J* = 7.9 Hz, 2H), 7.78 (d, *J* = 8.8 Hz, 1H), 7.69 (d, *J* = 8.7 Hz, 1H), 7.64 (d, *J* = 8.0 Hz, 2H), 6.16 (s, 1H), 4.39 (s, 1H), 3.71 (s, 2H), 3.66 (s, 1H), 3.63 (s, 3H), 3.20 – 2.97 (m, 2H), 2.27 (d, *J* = 11.6 Hz, 1H), 2.13 – 1.96 (m, 1H), 1.27 (d, *J* = 6.5 Hz, 3H), 1.22 (d, *J* = 6.0 Hz, 3H), 0.98 (d, *J* = 6.7 Hz, 4H). **$^{13}$C NMR** (100 MHz, $CDCl_3$) $\delta$ 169.77, 162.30, 154.53, 148.27, 145.04, 138.69, 136.58, 134.73, 129.61, 128.72, 128.28, 124.39, 122.43, 117.30, 107.03, 58.25, 53.74, 52.43, 51.12, 29.80, 29.12, 14.37, 9.78, 9.17, 8.44. **HRMS** (ESI-TOF) m/z: $[M + Na]^+$calculated for Chemical Formula: $C_{28}H_{30}N_8O_3SNa$, 581.2054, Found, 581.2040.

**8-((2*R*,5*R*)-4-(4-((3-cyclopropyl-1H-1,2,4-triazol-1-yl)sulfonyl)benzyl)-2,5-dimethylpiperazin-1-yl)-5-methyl-6-oxo-5,6-dihydro-1,5-naphthyridine-2-carbonitrile (SPM-25):**

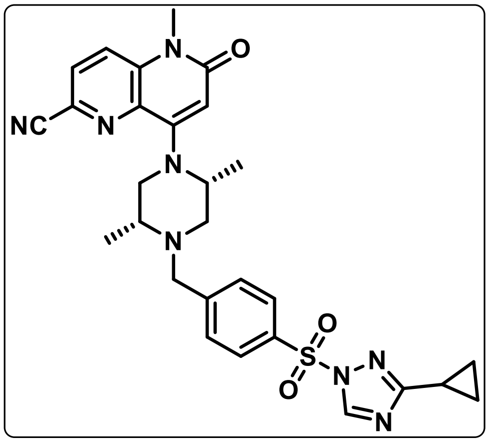


To a solution of **3c (**250 mg, 0.84 mmol), in anhydrous DCM (10 mL)**,** 4-((3-cyclopropyl-1H-1,2,4-triazol-1-yl) sulfonyl) benzaldehyde (175 mg, 0.63 mmol), sodium cyanoborohydride (150 mg, 2.4 mmol) were added. The resulting mixture was stirred at room temperature for 24 hrs, followed by addition of water (25 mL) and then extracted with ethyl acetate (3 × 25 mL). The organic layers were combined and dried over anhydrous $Na_2SO_4$. The residue was purified by Biotage flash chromatography using 95% EtOAc in *n*-hexane as an eluent to give the desired product **SPM-25** (0.080 g, 17 % yield) as a yellow solid. **$^1$H NMR** (400 MHz, $CDCl_3$) $\delta$ 8.57 – 8.50 (m, 1H), 8.02 (d, $J$ = 8.0 Hz, 2H), 7.78 (d, $J$ = 8.8 Hz, 1H), 7.70 (d, $J$ = 8.8 Hz, 2H), 7.63 (s, 1H), 6.15 (s, 1H), 4.85 (s, 1H), 4.18 (d, $J$ = 47.1 Hz, 1H), 3.63 (s, 3H), 3.44 (d, $J$ = 12.0 Hz, 1H), 3.20 (s, 2H), 2.59 (s, 2H), 2.11 – 1.98 (m, 2H), 1.23 (d, $J$ = 8.9 Hz, 6H), 0.97 (d, $J$ = 6.7 Hz, 4H). **$^{13}$C NMR** (100 MHz, $CDCl_3$) $\delta$ 169.74, 162.35, 154.56, 145.03, 138.58, 136.48, 134.71, 129.61, 128.70, 128.30, 124.45, 122.58, 117.23, 106.75, 60.48, 58.21, 53.71, 52.42, 51.01, 38.07, 29.17, 21.14, 14.33, 14.29, 9.69, 9.15, 8.42. **HRMS** (ESI-TOF) m/z: $[M + Na]^+$ calculated for Chemical Formula: $C_{28}H_{30}N_8O_3SNa$, 581.2054, Found, 581.2052

**8-((2*S*,5*S*)-4-(4-((3-cyclopropyl-1H-1,2,4-triazol-1-yl)sulfonyl)benzyl)-2,5-dimethylpiperazin-1-yl)-5-methyl-6-oxo-5,6-dihydro-1,5-naphthyridine-2-carbonitrile (SPM-26):**

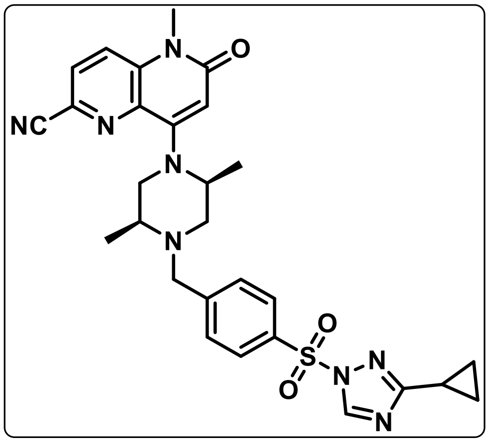

To a solution of **3d (**250 mg, 0.84 mmol), in anhydrous DCM (10 mL)**,** 4-((3-cyclopropyl-1H-1,2,4-triazol-1-yl) sulfonyl) benzaldehyde (175 mg, 0.63 mmol), sodium cyanoborohydride (150 mg, 2.4 mmol) were added. The resulting mixture was stirred at room temperature for 24 hrs, followed by addition of water (25 mL) and then extracted with ethyl acetate (3 × 25 mL). The organic layers were combined and dried over anhydrous $Na_2SO_4$. The residue was purified by Biotage flash chromatography using 95% EtOAc in *n*-hexane as an eluent to give the desired product **SPM-26** (0.088 g, 19% yield) as a yellow solid. **$^1$H NMR** (400 MHz, $CDCl_3$) $\delta$ 8.54 (s, 1H), 8.01 (d, *J* = 8.0 Hz, 2H), 7.81 – 7.75 (m, 1H), 7.69 (d, *J* = 8.8 Hz, 1H), 7.62 (d, *J* = 8.0 Hz, 2H), 6.14 (s, 1H), 4.84 (s, 1H), 4.23 (d, *J* = 14.7 Hz, 1H), 3.63 (s, 4H), 3.42 (d, *J* = 12.9 Hz, 1H), 3.18 (dd, *J* = 13.4, 10.0 Hz, 2H), 2.71 (s, 2H), 2.57 (s, 2H), 2.03 (d, *J* = 5.5 Hz, 1H), 1.24 – 1.17 (m, 6H), 0.97 (d, *J* = 6.7 Hz, 4H). **$^{13}$C NMR** (100 MHz, $CDCl_3$) $\delta$ 169.78, 162.26, 154.52, 148.29, 145.04, 138.70, 136.59, 134.69, 129.63, 128.74, 128.28, 127.98, 124.36, 122.40, 117.34, 60.54, 58.26, 53.75, 52.41, 51.16, 38.12, 29.12, 21.20, 14.36, 14.32, 9.82, 9.19, 8.47. **HRMS** (ESI-TOF) m/z: $[M + Na]^+$ calculated for Chemical Formula: $C_{28}H_{30}N_8O_3SNa$, 581.2054, Found, 581.2054

**4-(((2*R*,5*S*)-4-(6-cyano-1-methyl-2-oxo-1,2-dihydro-1,5-naphthyridin-4-yl)-2,5-dimethylpiperazin-1-yl)methyl)benzenesulfonamide**

**(SV-2089):**

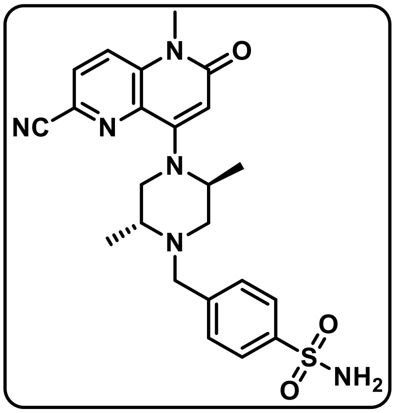

To a solution of **2a** (100 mg, 0.336 mol) in DCM (2mL), was adeed 4-formylbenzenesulfonamide (93 mg, 0.505 mol), and sodium cyanoborohydride (21 mg, 1mmol) at room temperature. The resulting mixture was stirred at room temperature for 24 hrs, followed by addition of water (50 mL) and then extracted with ethyl acetate (3 × 50 mL). The organic layers were combined and dried over anhydrous $Na_2SO_4$. The residue was purified by Biotage flash chromatography using 90% EtOAc in *n*-hexane as an eluent to give the desired product **SV2089** (0.063 g, 40% yield) as a white solid. **$^1$H NMR** (400 MHz, $CDCl_3$) $\delta$ 7.89 (d, $J$ = 8.0 Hz, 2H), 7.78 (d, $J$ = 8.7 Hz, 1H), 7.70 (d, $J$ = 8.8 Hz, 1H), 7.55 (d, $J$ = 8.0 Hz, 2H), 6.16 (s, 1H), 5.22 (s, 2H), 4.45 – 4.32 (m, 1H), 3.67 (d, $J$ = 15.9 Hz, 4H), 3.62 (s, 3H), 3.15 – 3.08 (m, 1H), 3.04 (dd, $J$ = 11.6, 3.6 Hz, 1H), 2.28 (dd, $J$ = 11.6, 3.0 Hz, 1H), 1.26 (d, $J$ = 6.9 Hz, 3H), 1.22 (d, $J$ = 6.5 Hz, 3H). **$^{13}$C NMR** (100 MHz, $CDCl_3$) $\delta$ 162.18, 154.37, 144.65, 140.66, 138.46, 136.37, 129.08, 128.08, 126.38, 124.13, 122.28, 117.15, 106.72, 58.05, 53.45, 52.24, 50.99, 50.78, 28.94, 14.20. **HRMS** (ESI-TOF) *m/z*: [M + Na]$^+$ calculated for $C_{23}H_{27}N_6O_3SNa$ 489.1679, Found, 489.1685.

**4-((2*R*,5*S*)-4-(6-cyano-1-methyl-2-oxo-1,2-dihydro-1,5-naphthyridin-4-yl)-2,5-dimethylpiperazin-1-yl)methyl)benzenesulfonic acid**

**(SV-2094):**

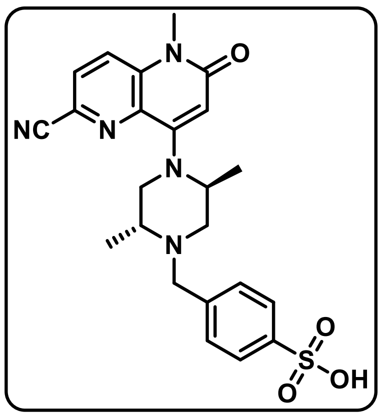

To a stirred solution of **AHL-7160** (100 mg, 0.179 mmol) in (2 mL) ACN–$H_2O$ (4:1), formic acid (20 μL) was added, and the reaction mixture was heated at 65 °C for 16 hrs. The reaction mixture was then cooled to room temperature and concentrated on a rotary evaporator. The remaining white residue was washed with DCM (1 mL × 2) and dried in vacuo to afford **SV2094** (0.082 g, 95% yield) as a white solid. **$^1$H NMR** (400 MHz, $CD_3OD$) *δ* 8.03 (tdd, *J* = 17.2, 8.8, 2.0 Hz, 4H), 7.81 (d, *J* = 8.0 Hz, 2H), 6.15 (d, *J* = 1.9 Hz, 1H), 4.48 (s, 1H), 3.82 (d, *J* = 4.2 Hz, 2H), 3.72 (s, 2H), 3.65 (d, *J* = 1.8 Hz, 3H), 3.18 (s, 1H), 3.11 (dt, *J* = 12.1, 2.6 Hz, 1H), 2.34 (dd, *J* = 11.7, 3.1 Hz, 1H), 1.30 (dd, *J* = 6.6, 1.8 Hz, 3H), 1.26 (dd, *J* = 6.6, 1.8 Hz, 3H). **$^{13}$C NMR** (100 MHz, $CD_3OD$) *δ* 164.40, 156.53, 150.50, 139.92, 137.28, 131.11, 129.84, 129.57, 125.93, 125.30, 118.18, 106.75, 59.03, 55.15, 53.83, 52.13, 51.97, 29.57, 14.60, 10.03. **HRMS** (ESI-TOF) *m*/*z*: $[M + Na]^+$ calculated for $C_{23}H_{26}N_6O_3SNa$ 490.1526, Found, 490.1521.

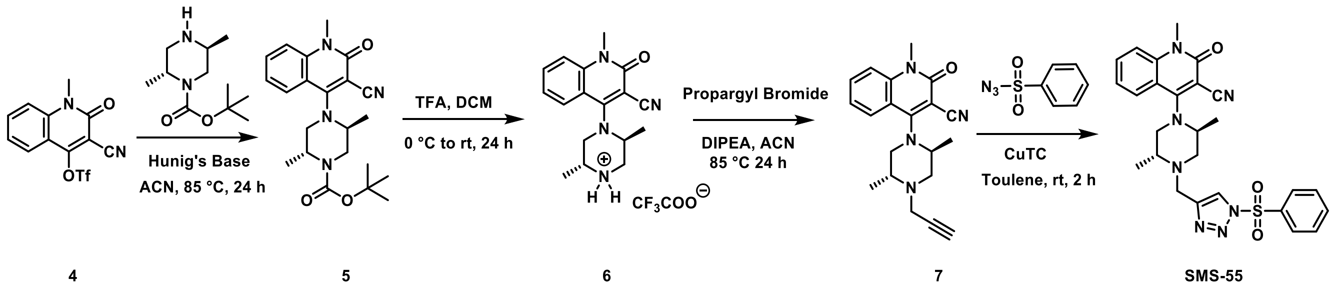


***tert*-butyl (2*R*,5*S*)-4-(3-cyano-1-methyl-2-oxo-1,2-dihydroquinolin-4-yl)-2,5-dimethylpiperazine-1-carboxylate (5):**

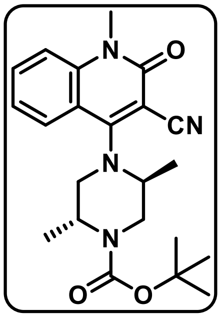


To a stirred solution of **4** (2.2 g, 6.62 mmol) and commercially available *tert*-butyl (2*R*, 5*S*)-2,5-dimethylpiperyzine-1-caroboxylate (1.4 g, 6.62 mmol) in anhydrous acetonitrile (15 mL), Hunig's base (1.8 mL, 13.24 mmol) was added at room temperature. The reaction mixture was stirred at 85 °C for 16 hrs, after which it was cooled down to room temperature and diluted with brine (30 mL) and then extracted with $Et_2O$ (2 × 30 mL). The combined organic layer was dried over $Na_2SO_4$. The residue was purified by Biotage flash chromatography using a gradient of 0% to 40% EtOAc in *n*-hexane as an eluent to give the desired product **5** (2.5 g, 96% yield) as a yellow solid. **$^1$H NMR** (600 MHz, $CDCl_3$) $\delta$ 7.83 (d, *J* = 8.1 Hz, 1H), 7.66 (ddd, *J* = 8.6, 7.1, 1.5 Hz, 1H), 7.39 (dd, *J* = 8.6, 1.0 Hz, 1H), 7.30 – 7.26 (m, 1H), 4.49 (s, 1H), 4.33 (dd, *J* = 12.5, 3.9 Hz, 1H), 4.16 – 4.08 (m, 1H), 3.87 (d, *J* = 13.3 Hz, 1H), 3.79 – 3.69 (m, 1H), 3.68 (s, 3H), 3.06 (d, *J* = 12.3 Hz, 1H), 1.49 (s, 9H), 1.28 (t, *J* = 6.6 Hz, 6H).

**(2*R*,5*S*)-4-(3-cyano-1-methyl-2-oxo-1,2-dihydroquinolin-4-yl)-2,5-dimethylpiperazin-1-ium-trifluoroacetate (6):**

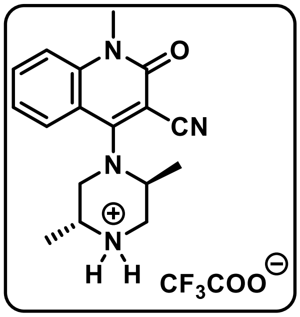


The compound **5** (2.5 g, 6.30 mmol) was dissolved in anhydrous dichloromethane (10 mL), and trifluoroacetic acid (5.1 mL, 63.09 mmol) was dropwise added at 0 °C under nitrogen ($N_2$) atmosphere for 10-15 min. The reaction mixture was stirred at room temperature for 16 hrs and then concentrated under reduced pressure. The resultant residue was washed in *n*-Pentane (3 x 20 mL) and $Et_2O$ (1 x 7 mL) and then dried under high vacuum to afford the desired final product **6** (2.8 g, 100% yield) as a white solid. **$^1$H NMR** (600 MHz, $CDCl_3$) $\delta$ 8.14 (s, 1H), 7.80 (t, *J* = 7.8 Hz, 1H), 7.57 – 7.37 (m, 2H), 4.38 (s, 1H), 3.85 (s, 1H), 3.78 (s, 3H),

3.62 (d, $J$ = 51.3 Hz, 2H), 3.27 (d, $J$ = 87.9 Hz, 2H), 1.46 (d, $J$ = 5.4 Hz, 3H), 1.12 (d, $J$ = 5.8 Hz, 3H).

**4-((2*S*,5*R*)-2,5-dimethyl-4-(prop-2-yn-1-yl)piperazin-1-yl)-1-methyl-2-oxo-1,2-dihydroquinoline-3-carbonitrile (7):**

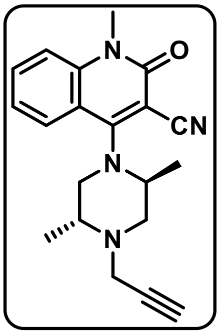


The compound **6** (2.8 g, 9.45 mmol) and propargyl bromide (1 mL, 11.81 mmol) were dissolved in anhydrous acetonitrile (30 mL), and *N, N*-Diisopropylethylamine base (5 mL, 28.35 mmol) was added at room temperature. The reaction mixture was stirred at 85 °C for 16 hrs, after which it was diluted with water (30 mL) and then extracted with ethyl acetate (30 mL), and the aqueous layer was separated and ethyl acetate (3 × 30 mL). The combined organic layer was dried over $Na_2SO_4$. The residue was purified by Biotage flash chromatography using a gradient of 0% to 65% EtOAc in *n*-hexane as an eluent to give the desired product **7** (1.4 g, 45% yield) as a yellow solid. **$^1$H NMR** (600 MHz, $CDCl_3$) $\delta$ 8.14 (d, $J$ = 8.1 Hz, 1H), 7.69 (ddd, $J$ = 8.6, 7.2, 1.6 Hz, 1H), 7.38 (dd, $J$ = 8.6, 1.0 Hz, 1H), 7.31 (ddd, $J$ = 8.1, 7.1, 1.0 Hz, 1H), 4.20 (s, 1H), 3.72 (s, 3H), 3.70 (s, 1H), 3.43 (dd, $J$ = 17.5, 2.4 Hz, 1H), 3.19 (s, 1H), 3.06 – 2.96 (m, 1H), 2.93 – 2.82 (m, 2H), 2.60 (dd, $J$ = 11.3, 9.4 Hz, 1H), 2.31 (t, $J$ = 2.4 Hz, 1H), 1.06 (dd, $J$ = 7.1, 6.2 Hz, 6H).

**4-((2*S*,5*R*)-2,5-dimethyl-4-((1-(phenylsulfonyl)-1H-1,2,3-triazol-4-yl)methyl)piperazin-1-yl)-1-methyl-2-oxo-1,2-dihydroquinoline-3-carbonitrile**

**(SMS-55):**

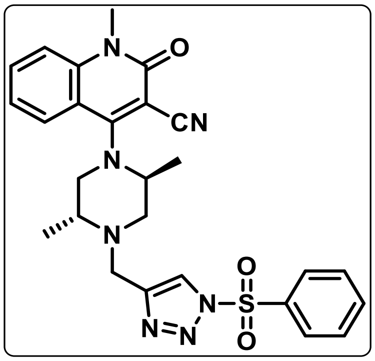

To a stirred solution of **7** (200 mg, 0.598 mmol) in anhydrous toluene (5 mL), benzene sulfonyl azide (160 mg, 0.718 mmol), Copper (ι)-thiophene-2-carboxylate (25 mg, 0.150 mmol) were added. The resulting mixture was stirred at room temperature for 2 hrs, followed by addition of water (30 mL) and then extracted with ethyl acetate (3 × 30 mL). The organic layers were combined and dried over anhydrous $Na_2SO_4$. The residue was purified by Biotage flash chromatography using a gradient of 0% to 90% EtOAc in *n*-hexane as an eluent to give the desired product **SMS-55** (0.300 g, 97% yield) as a yellow solid. **$^1$H NMR** (400 MHz, $CDCl_3$) $\delta$ 8.19 – 8.14 (m, 2H), 8.12 (s, 1H), 8.00 (d, $J$ = 8.2 Hz, 1H), 7.79 – 7.71 (m, 1H), 7.71 – 7.66 (m, 1H), 7.66 – 7.58 (m, 2H), 7.38 (d, $J$ = 8.5 Hz, 1H), 7.32 – 7.27 (m, 1H), 4.12 (s, 1H), 3.98 (d, $J$ = 14.9 Hz, 1H), 3.86 (d, $J$ = 14.9 Hz, 1H), 3.71 (s, 3H), 3.02 (t, $J$ = 11.4 Hz, 2H), 2.80 (s, 1H), 2.36 – 2.21 (m, 1H), 1.76 (s, 1H), 1.15 (d, $J$ = 6.2 Hz, 3H), 1.05 (d, $J$ = 6.2 Hz, 3H). **$^{13}$C NMR** (151 MHz, $CDCl_3$) $\delta$ 163.28, 160.34, 144.78, 141.23, 136.23, 135.81, 133.81, 129.98, 128.73, 126.48, 122.86, 122.50, 119.39, 115.44, 115.08, 60.47, 54.62, 48.42, 30.13, 21.15, 16.76, 14.30. **HRMS** (ESI-TOF) *m/z*: $[M + Na]^+$ calculated for $C_{26}H_{27}N_7O_3SNa$ 540.1788, Found, 540.1783.

### 4. NMR SPECTRA

$^1H$ NMR ($CDCl_3$ 600 MHz) of **2a**

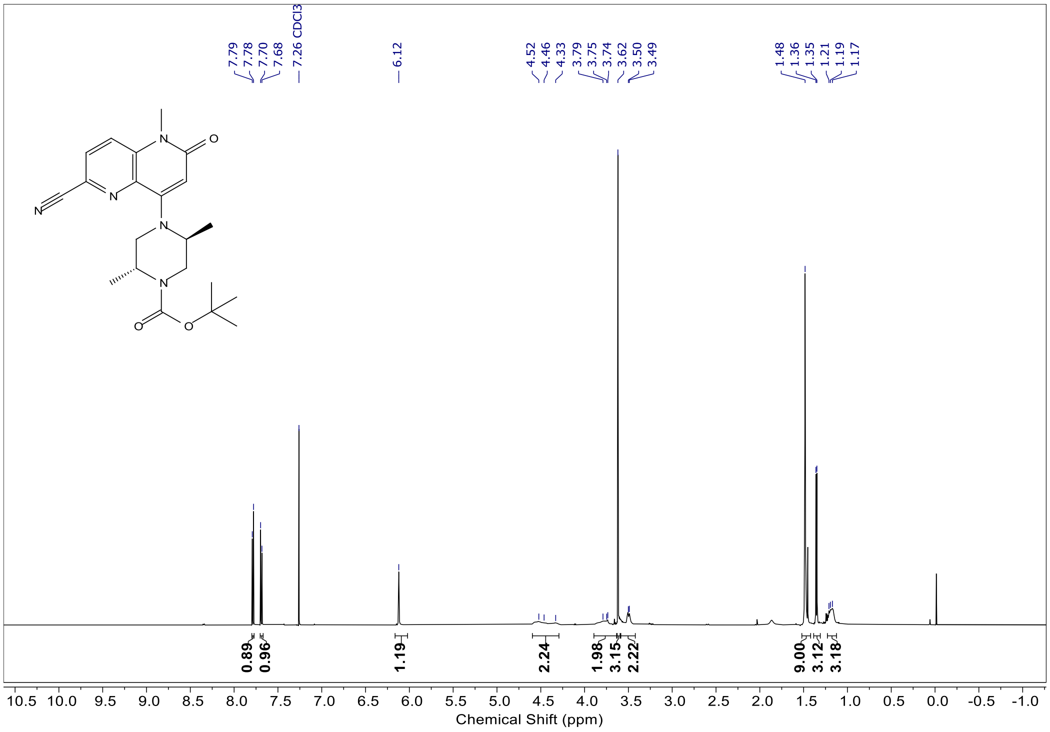

$^1$H NMR ($CDCl_3$ 400 MHz) of **2b**

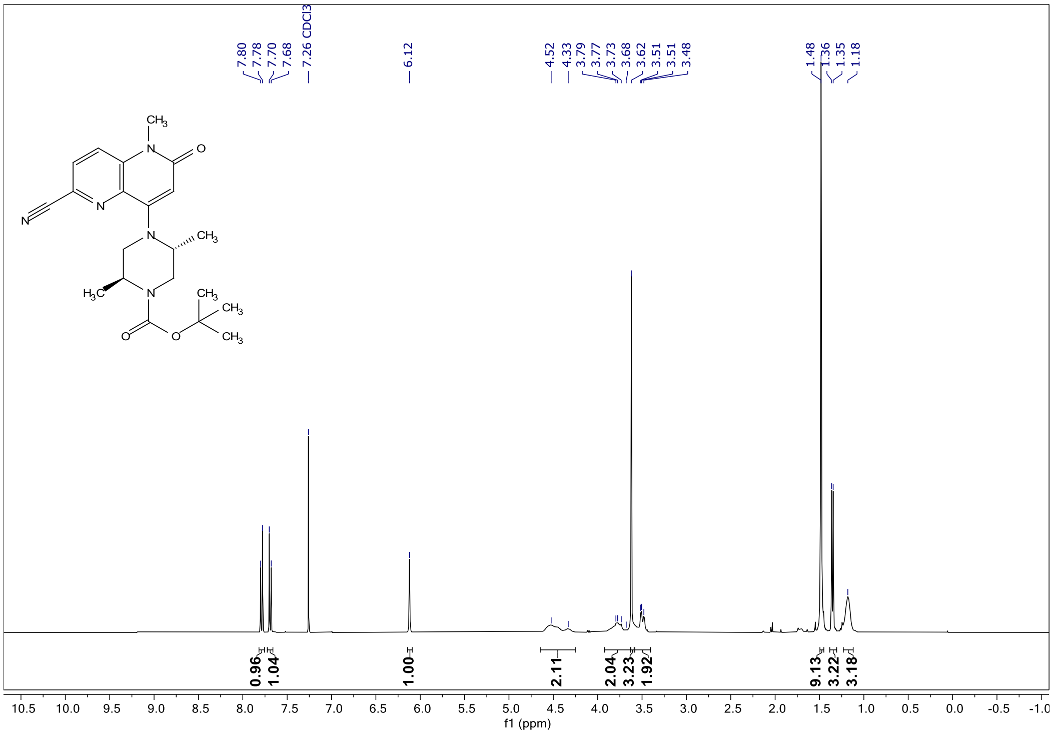

$^{1}$H NMR ($CDCl_3$ 400 MHz) of **2c**

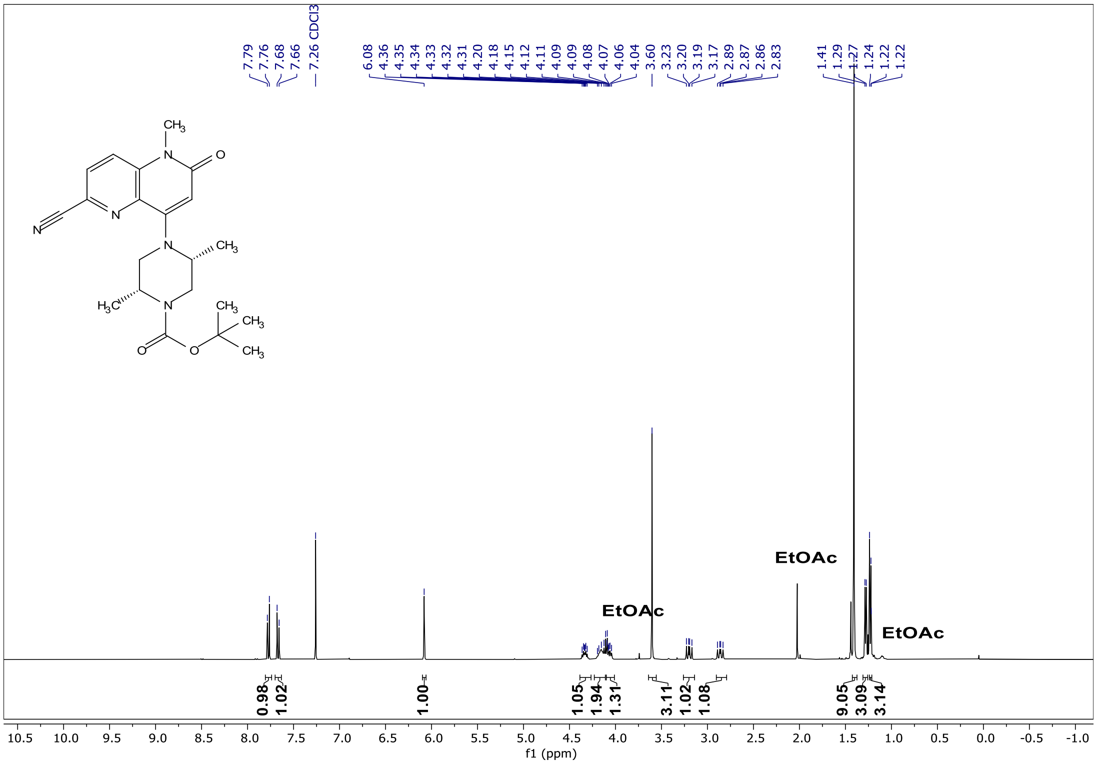

$^{1}$H NMR ($CDCl_3$ 400 MHz) of **2d**

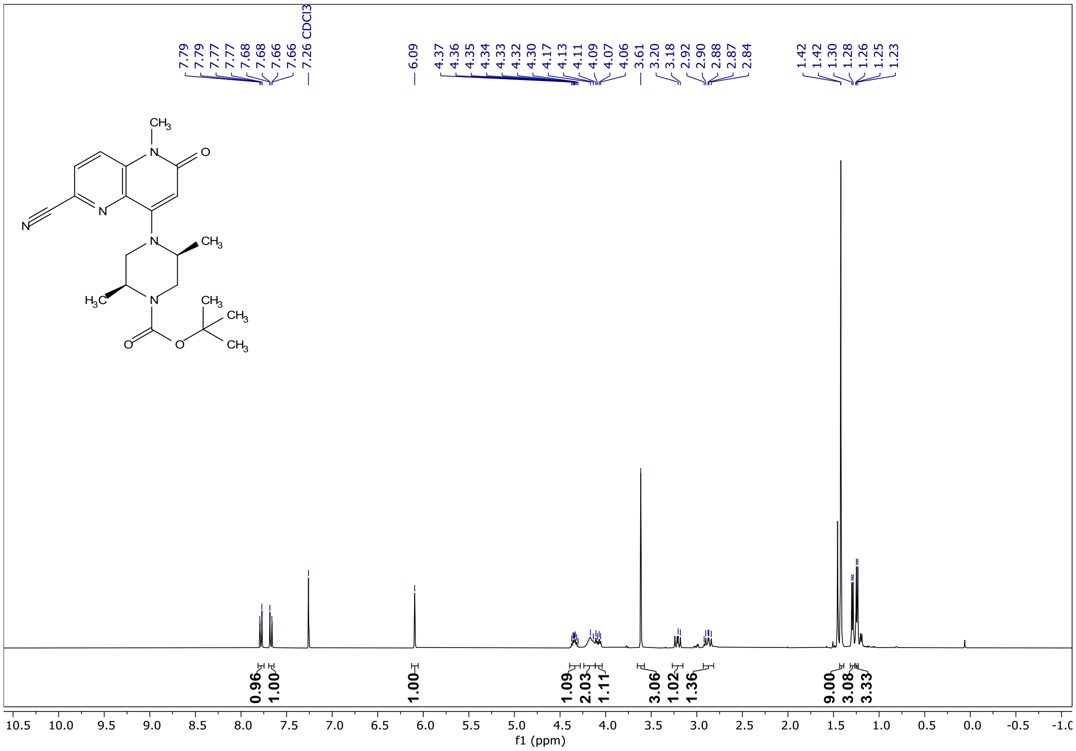

$^{1}H$ NMR ($CDCl_3$ 400 MHz) of **3a**

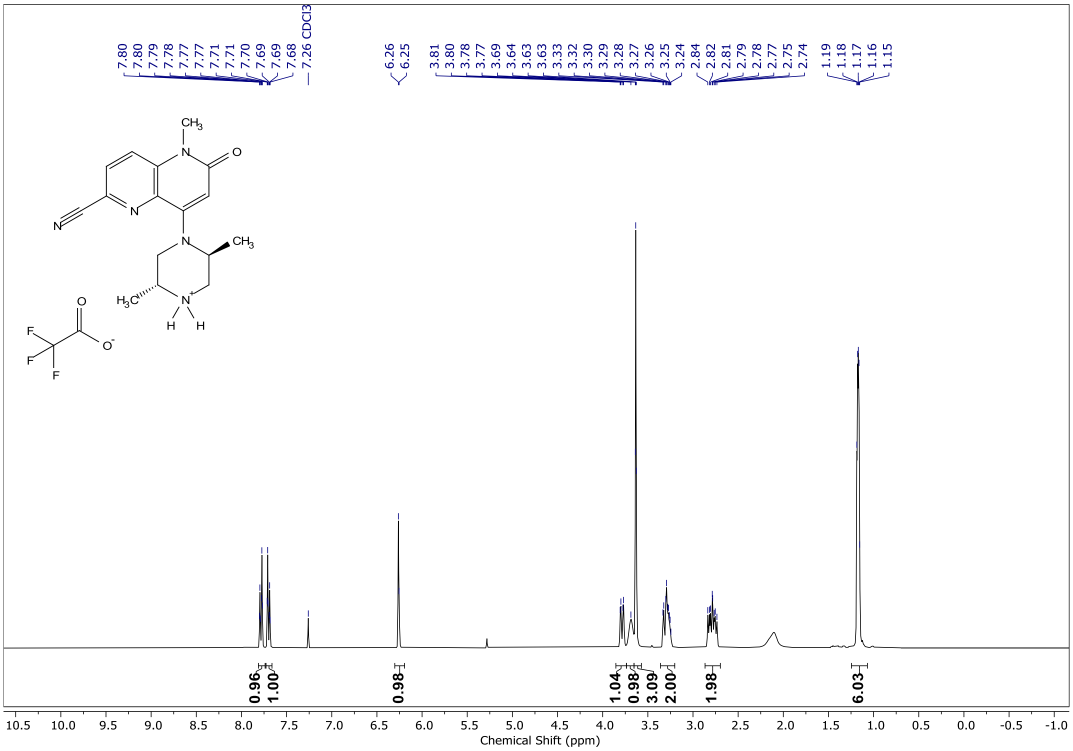

$^{1}$H NMR ($CDCl_3$ 400 MHz) of **3b**

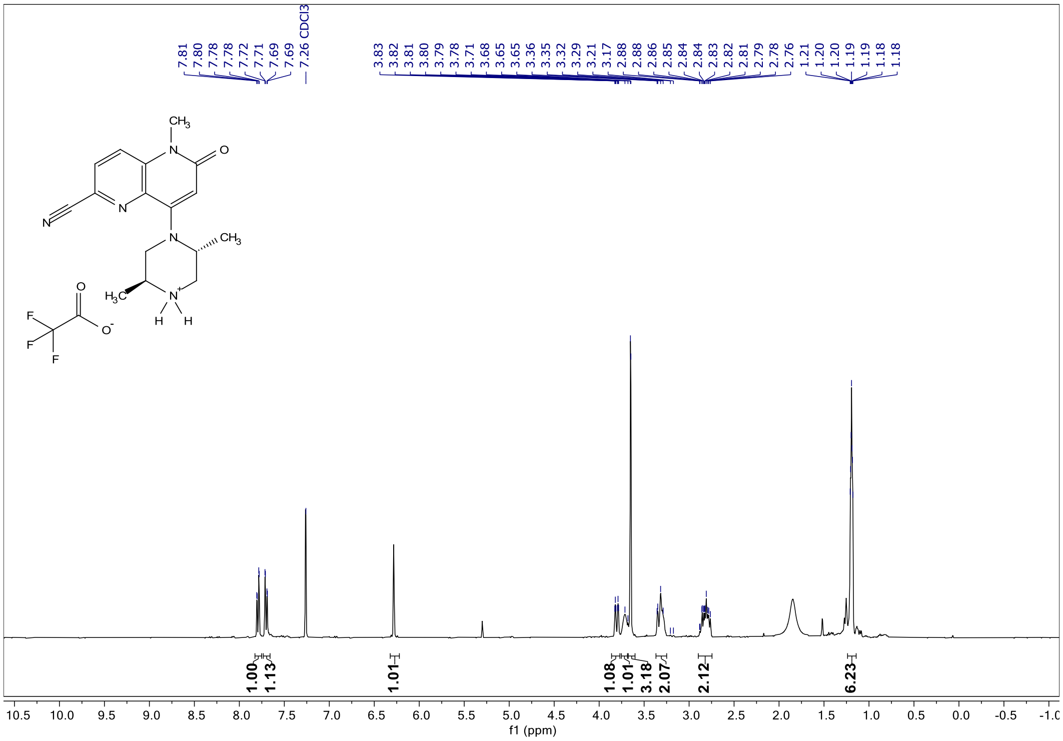

[1]H NMR (CDCl3 400 MHz) of **3c**

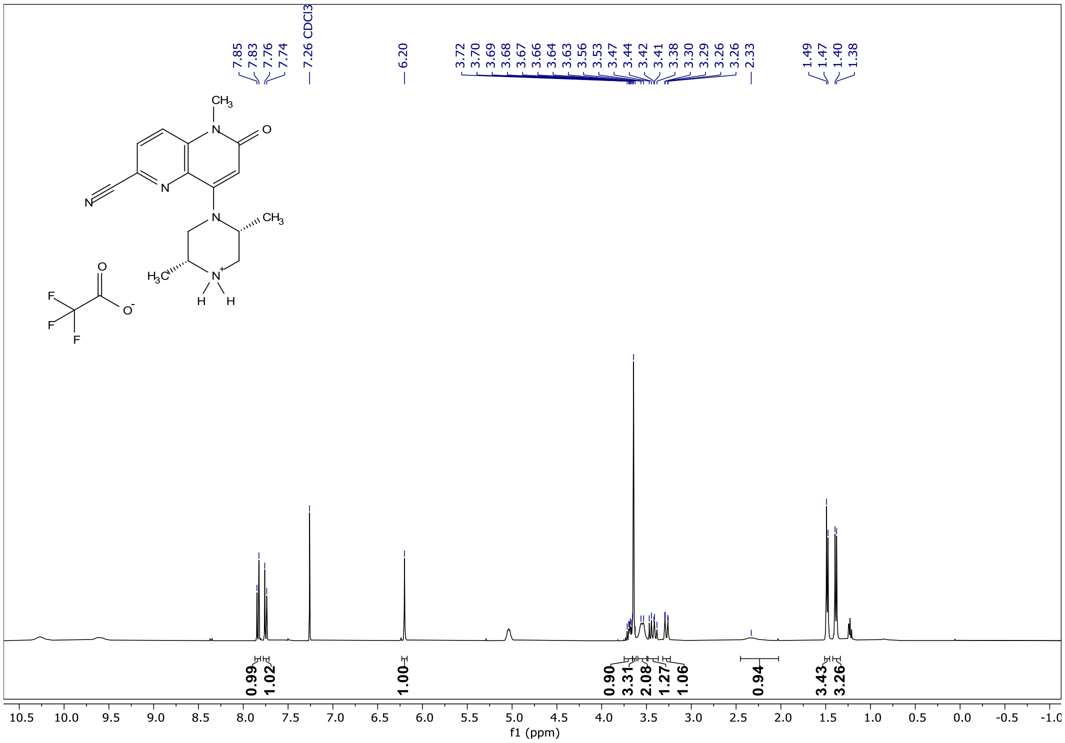

[1]H NMR ($CDCl_3$ 400 MHz) of **3d**

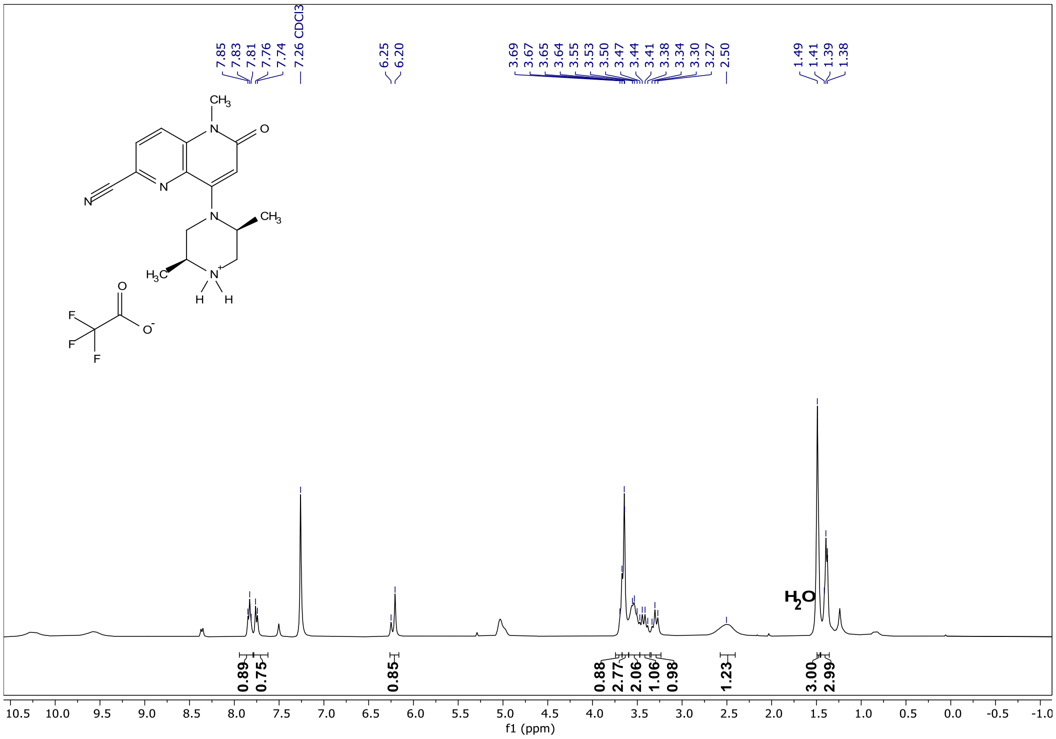

$^{1}$H NMR ($CDCl_3$ 600 MHz) of **4-((3-cyclopropyl-1H-1,2,4-triazol-1-yl)sulfonyl)benzaldehyde**

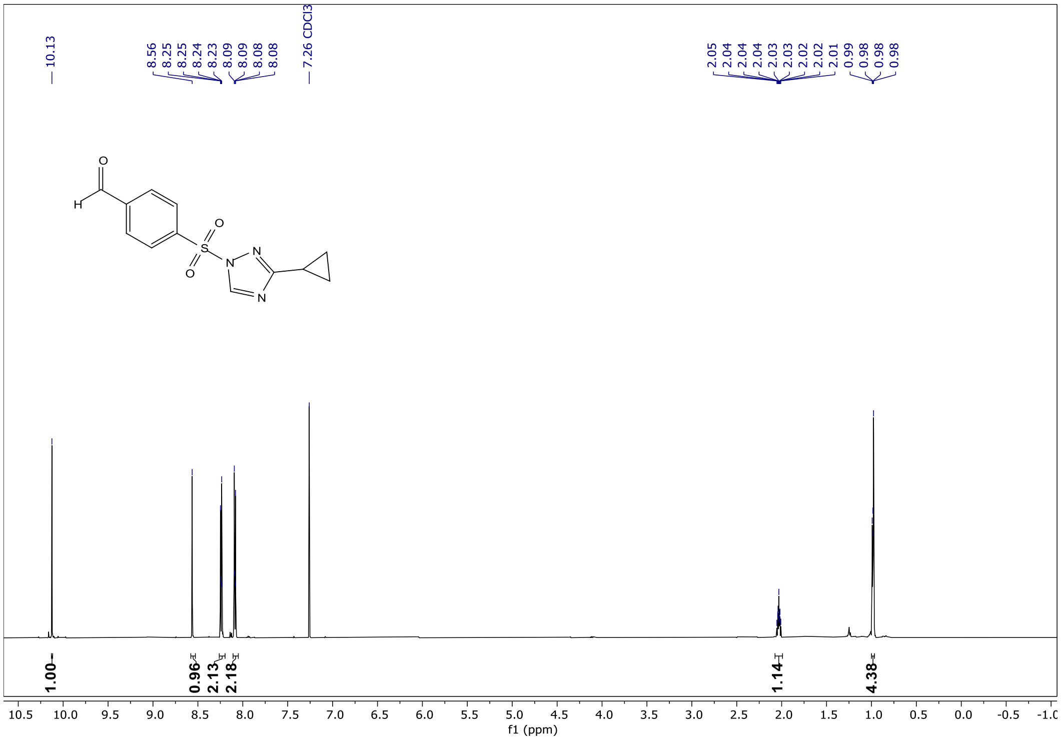

$^{1}H$ NMR ($CDCl_3$ 600 MHz) of **AHL-7160**

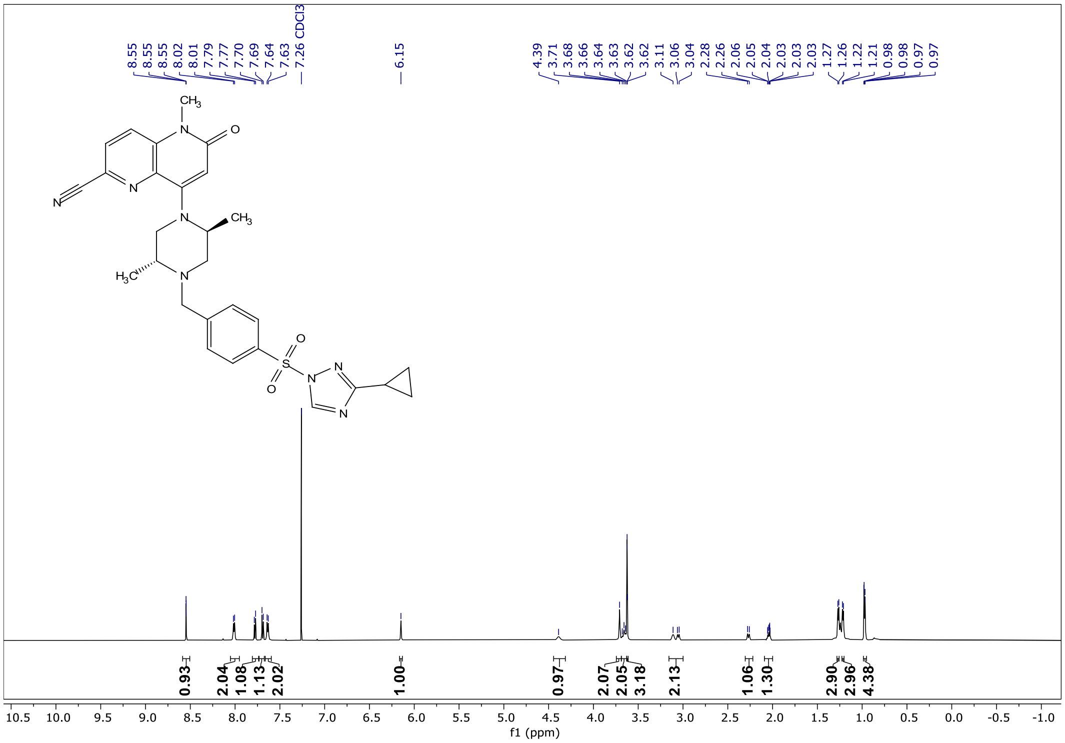

$^{13}$C NMR (CDCl$_3$ 100 MHz) of **AHL-7160**

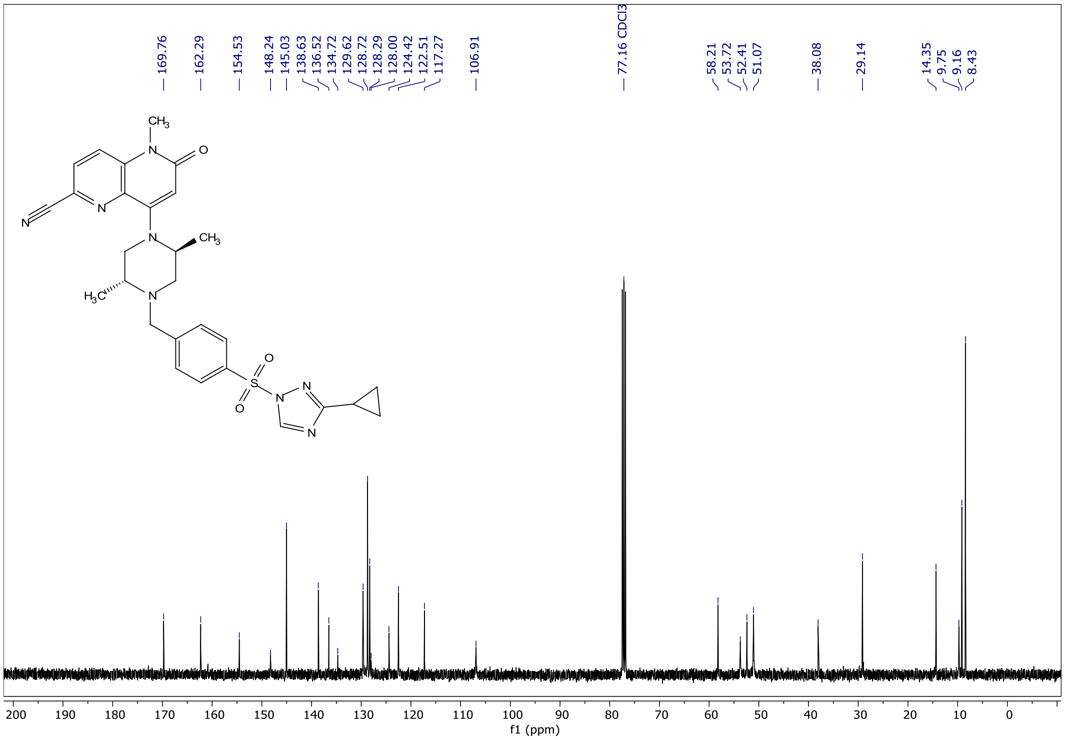

<sup></sup>[1]H NMR ($CDCl_3$ 400 MHz) of **SPM-04**

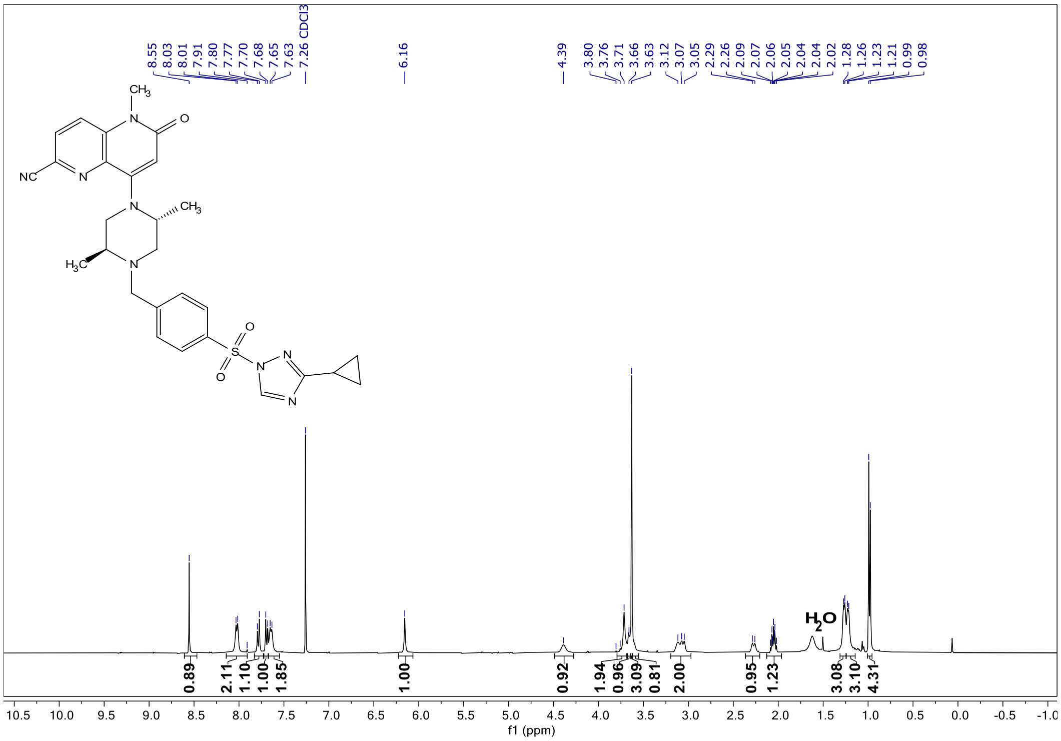

$^{13}C$ NMR ($CDCl_3$ 100 MHz) of **SPM-04**

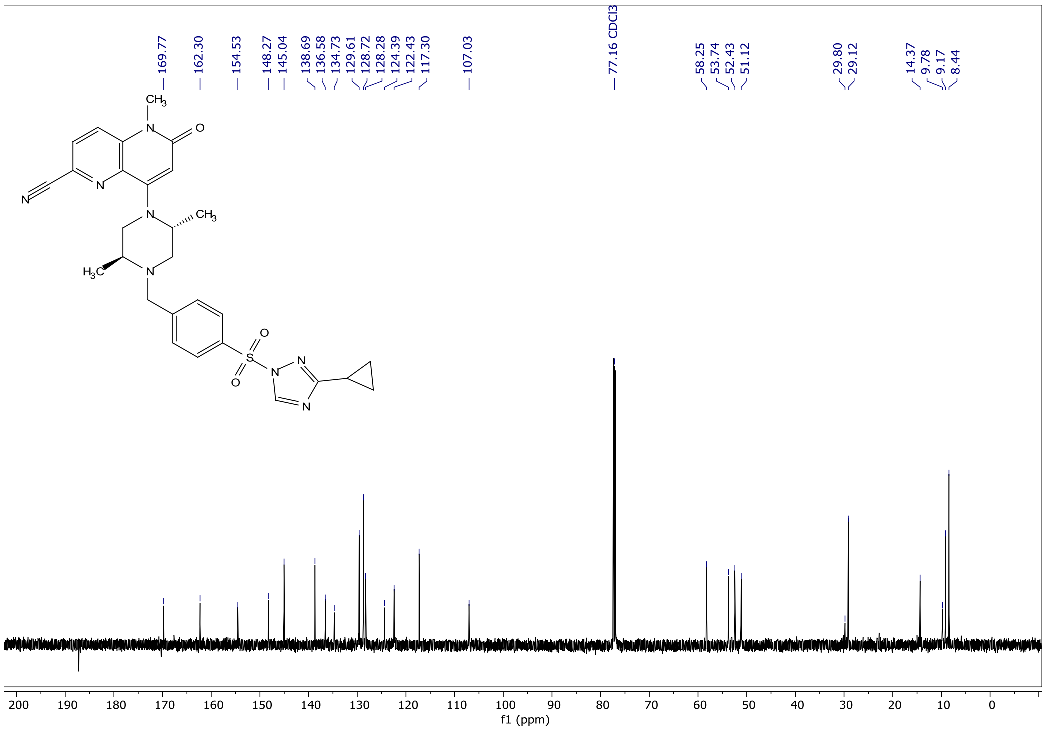

$^{1}H$ NMR ($CDCl_3$ 400 MHz) of **SPM-25**

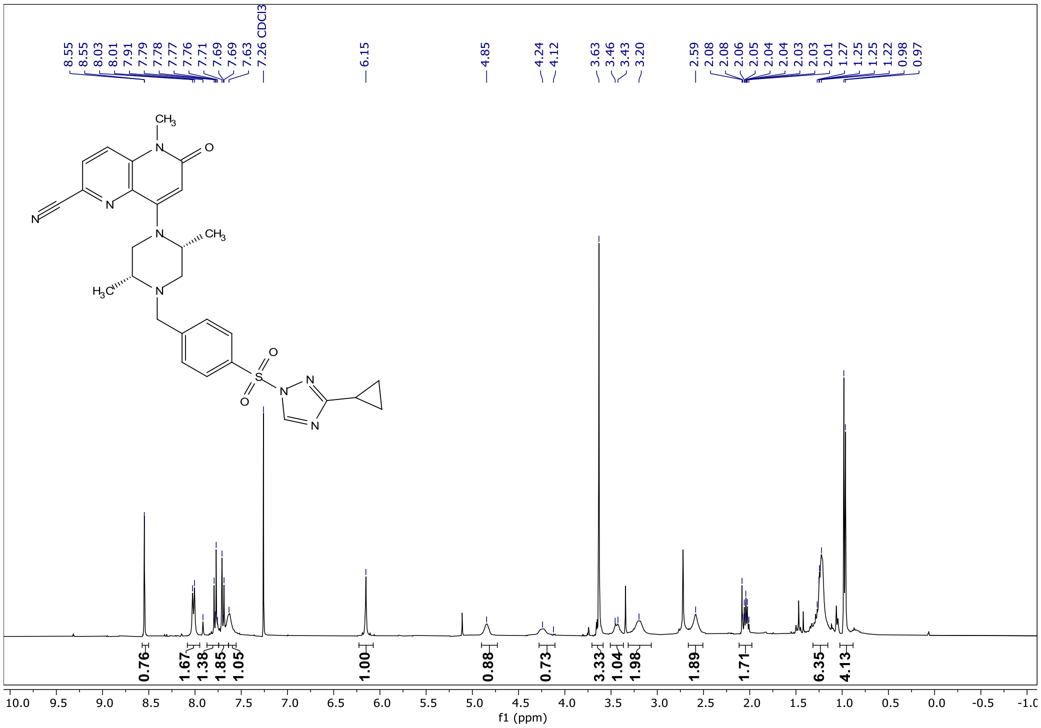

<sup></sup>

$^{13}$C NMR ($CDCl_3$ 100 MHz) of **SPM-25**

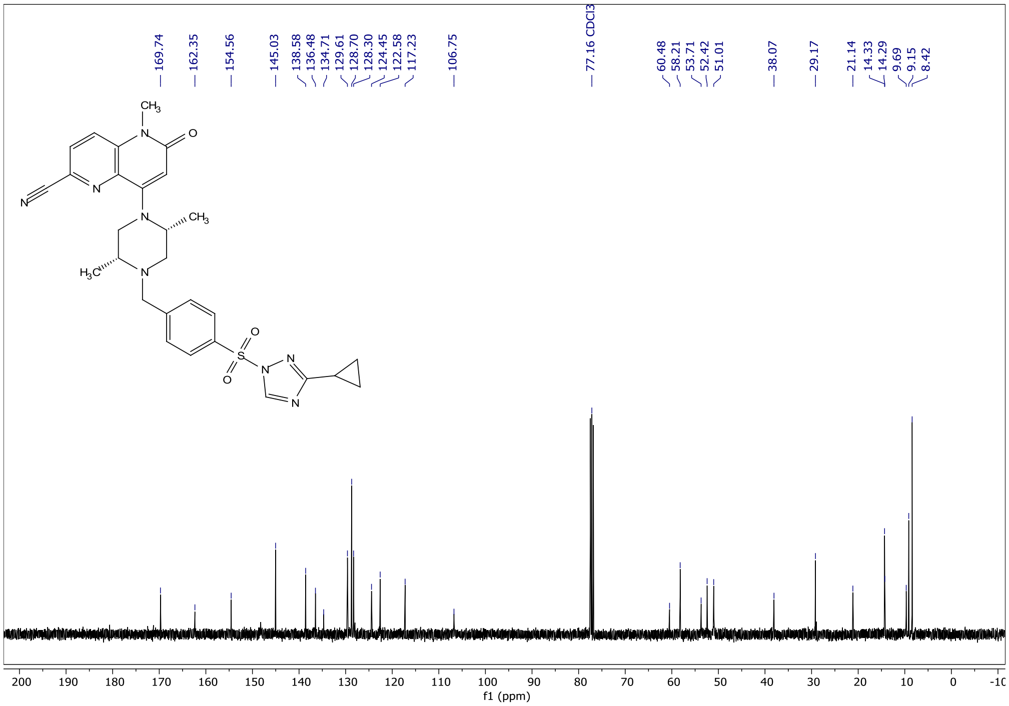

$^{1}$H NMR ($CDCl_3$ 400 MHz) of **SPM-26**

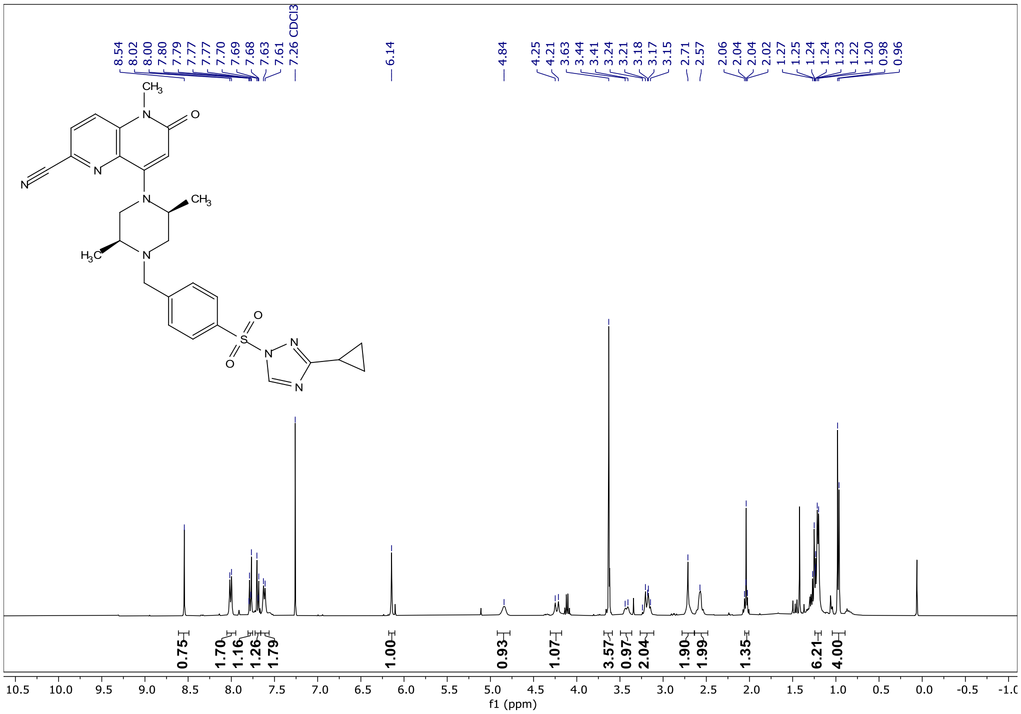

$^{13}C$ NMR ($CDCl_3$ 100 MHz) of **SPM-26**

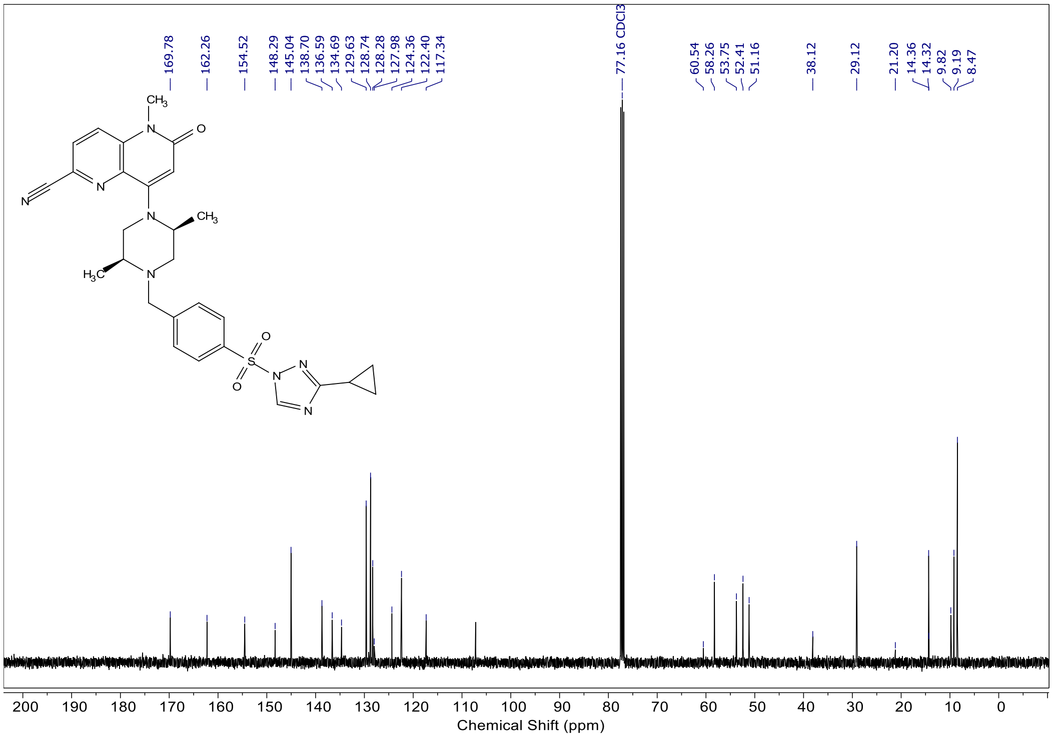

$^{1}H$ NMR ($CDCl_3$ 400 MHz) of **SV-2089**

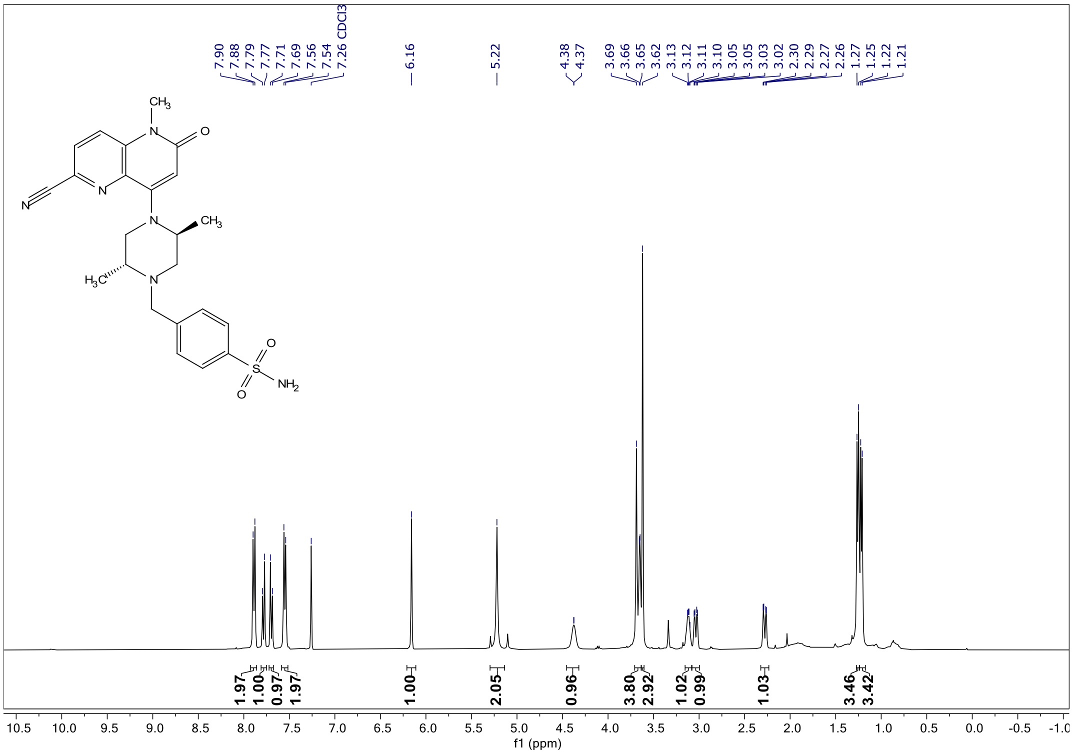

$^{13}C$ NMR ($CDCl_3$ 100 MHz) of **SV-2089**

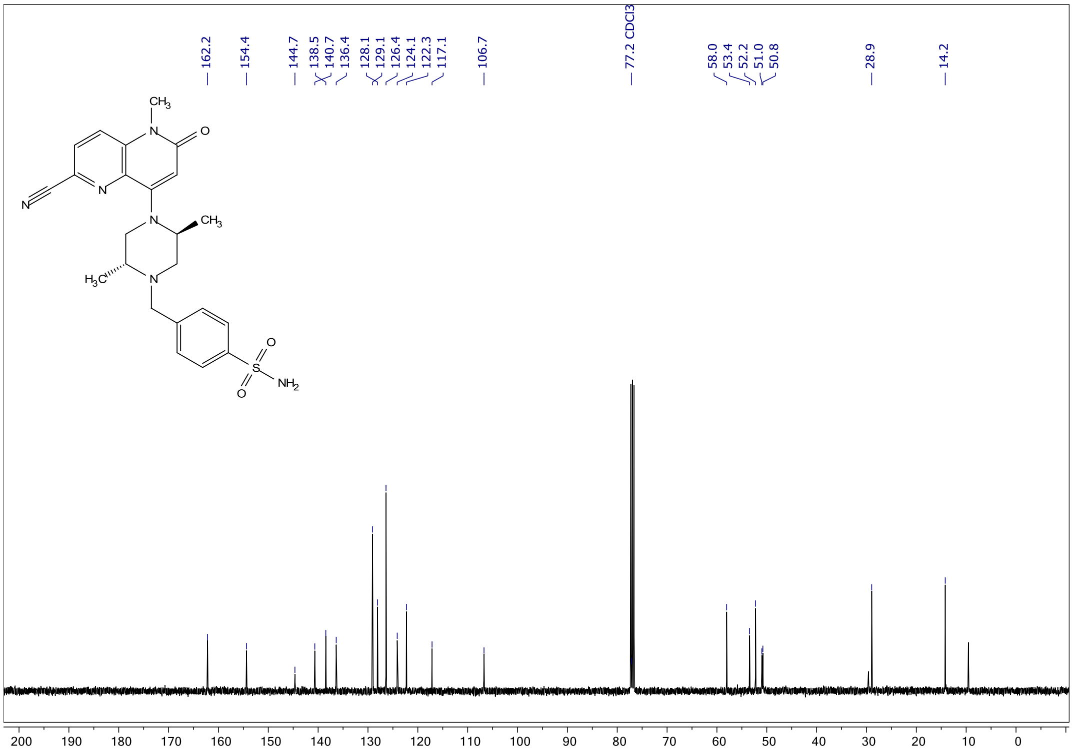

<sup></sup>$^{1}$H NMR ($CD_3OD$ 400 MHz) of **SV-2094**

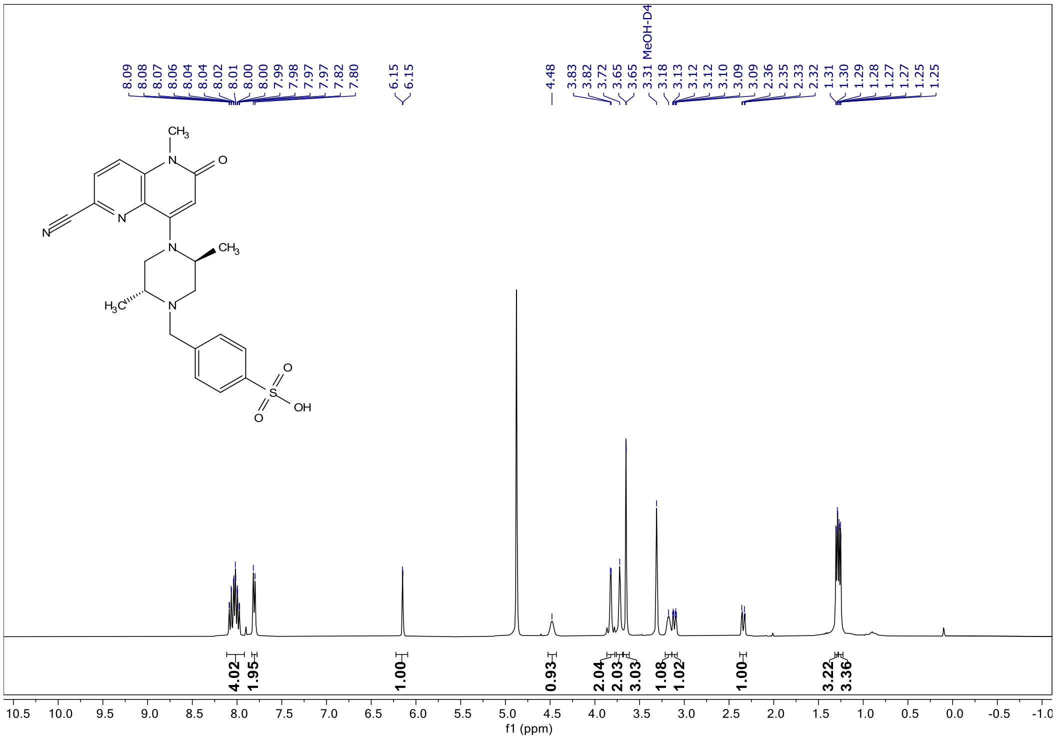

$^{13}$C NMR ($CD_3OD$ 100 MHz) of **SV-2094**

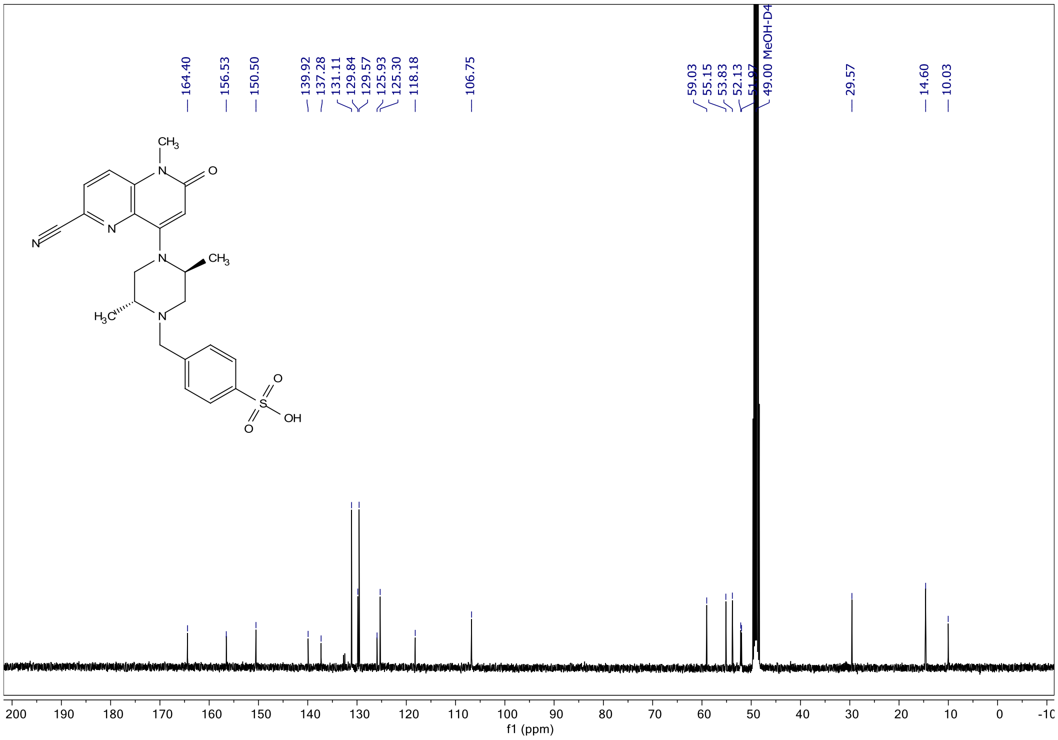

$^{1}$H NMR ($CDCl_3$ 600 MHz) of **5**

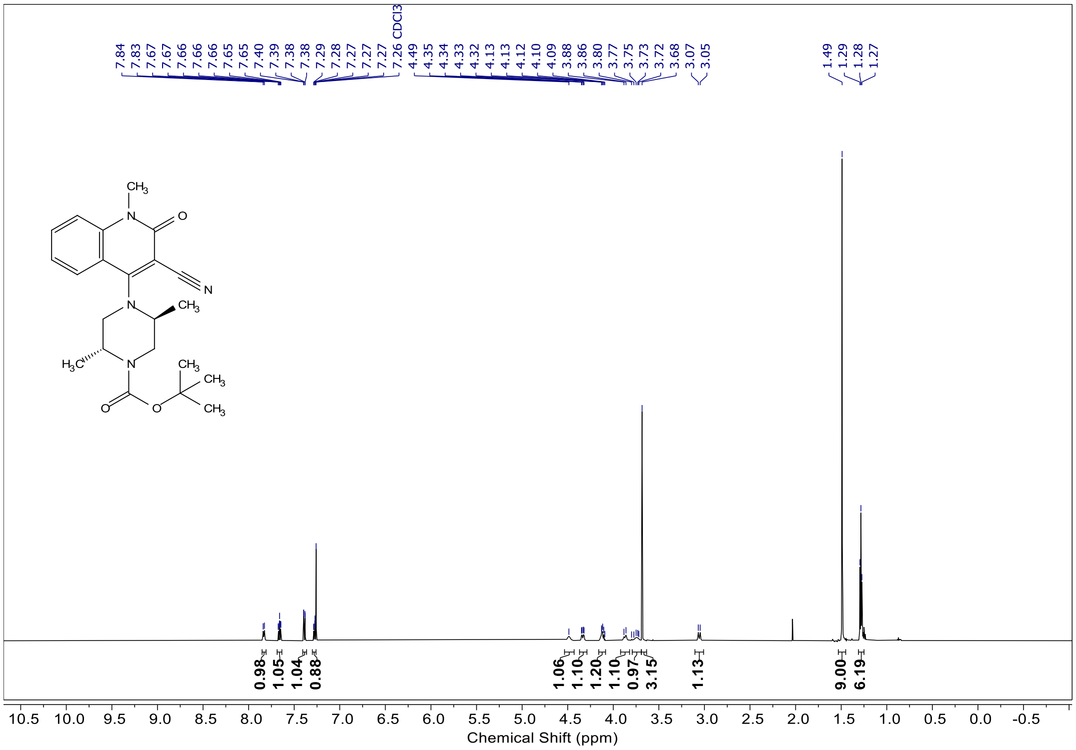

$^{1}$H NMR ($CDCl_3$ 600 MHz) of **6**

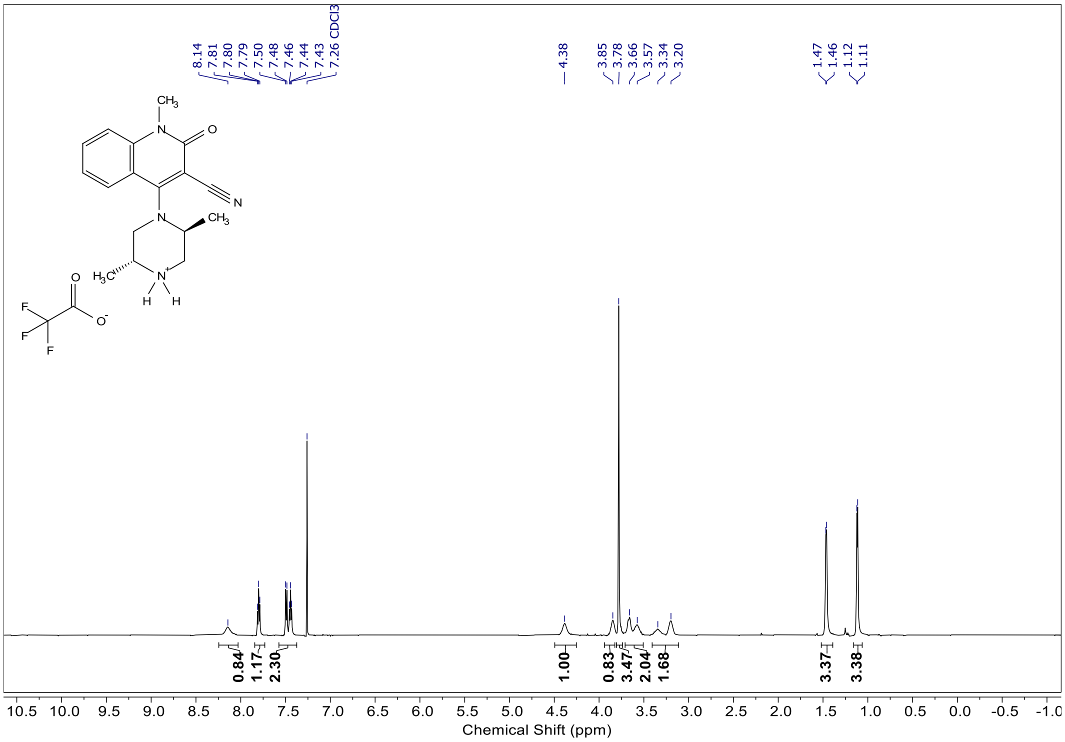

$^{1}$H NMR ($CDCl_3$ 600 MHz) of **7**

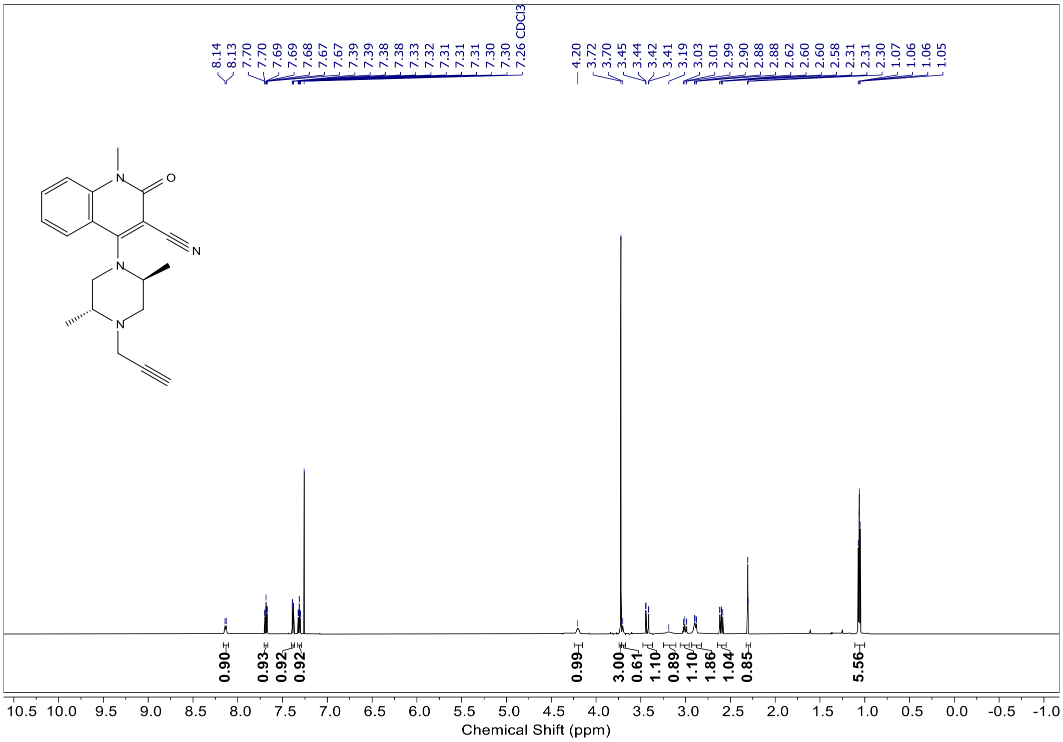

<sup></sup>$^{1}$H NMR ($CDCl_3$ 400 MHz) of **SMS-55**

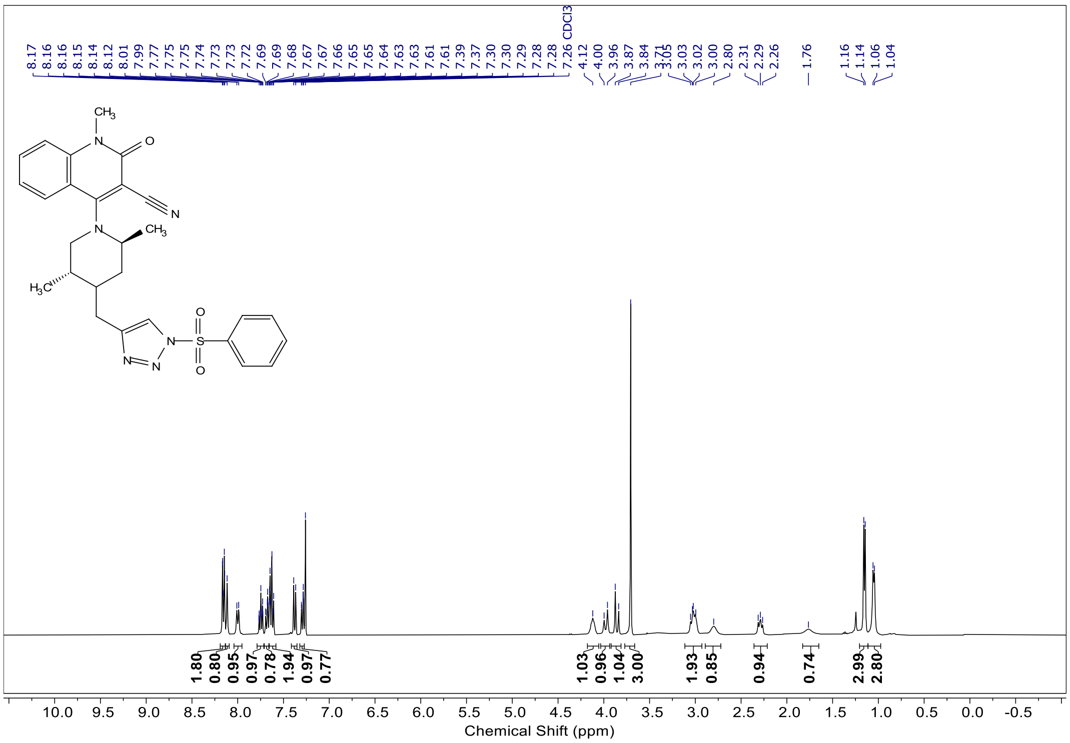

$^{13}$C NMR ($CDCl_3$ 151 MHz) of **SMS-55**

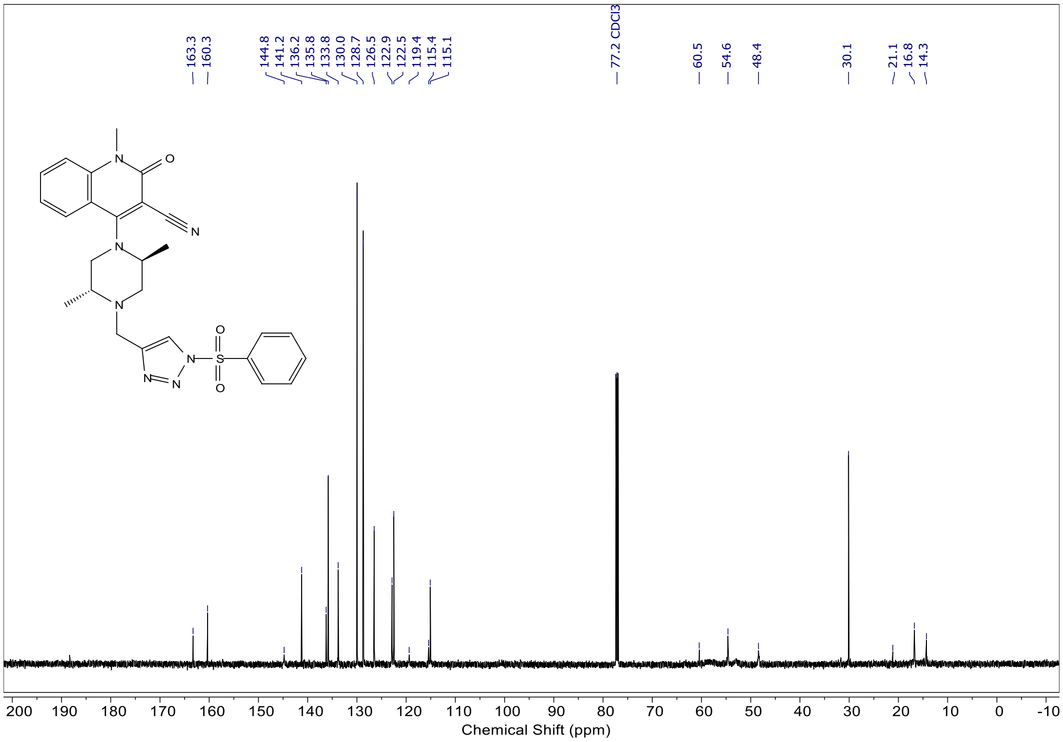
